# Phylogenetic endemism of the world’s seed plants

**DOI:** 10.1101/2022.12.28.522105

**Authors:** Lirong Cai, Holger Kreft, Amanda Taylor, Julian Schrader, Wayne Dawson, Franz Essl, Mark van Kleunen, Jan Pergl, Petr Pyšek, Marten Winter, Patrick Weigelt

## Abstract

Assessing phylogenetic endemism, i.e., the distribution of geographically restricted and evolutionarily unique species, is key to understanding biogeographic patterns and processes and critical for global conservation planning. Here, we quantified the geographic distribution and drivers of phylogenetic endemism for ~320,000 seed plants worldwide and identified centers and drivers of evolutionarily young (neoendemism) and evolutionarily old endemism (paleoendemism). Tropical and subtropical islands of the Southern Hemisphere as well as tropical mountainous regions displayed the world’s highest phylogenetic endemism. Tropical moist forests (e.g. Amazonia) and continental islands in south-east Asia emerged as centers of paleoendemism, while both high neo- and paleoendemism were found on ancient continental fragment islands (e.g. Madagascar) and in Mediterranean-climate regions. Global variation in phylogenetic endemism was best explained by a combination of past and present environmental factors (80.3% – 88.1% of variance explained). Geographic isolation and environmental heterogeneity emerged as primary drivers promoting high phylogenetic endemism. Also, warm and wet climates with long-term climatic stability showed a significant positive effect. However, environmental effects on phylogenetic endemism varied with geographic isolation, reflecting the unique evolutionary and biogeographic dynamics on oceanic islands. Long-term climatic stability promoted the persistence of paleoendemics, while isolation promoted higher neoendemism leading to oceanic islands and tropical mountainous regions being centers of both neo- and paleoendemism. Our study provides new insights into the evolutionary underpinnings of biogeographic patterns in seed plants, and by identifying areas of high evolutionary and biogeographic uniqueness, it may serve as a key resource for setting global conservation priorities.

## Introduction

Plant species range sizes vary widely from being nearly global to extremely small, e.g. being restricted to a single mountain or island (1). Understanding the global geographic distribution of range-restricted species (endemics) and the mechanisms that create areas of high endemism is a central question in biogeography with profound implications for biodiversity conservation (2). The small ranges of endemic species make them more likely to be threatened with extinction (3, 4). If endemic species are simultaneously evolutionarily unique, their loss leads to high losses of evolutionary history (5–7). Furthermore, evolutionarily unique endemics are likely to be associated with irreplaceable ecological and functional characteristics (8, 9). It is therefore important to account for the phylogenetic position and evolutionary uniqueness of species when assessing endemism.

Phylogenetic endemism (PE) is a measure that accounts for the phylogenetic uniqueness of range-restricted species (10). Regions with high PE harbor more evolutionarily unique lineages with restricted geographic distributions than regions with low PE. Assessing PE is thus crucial for understanding the evolutionary mechanisms underlying biodiversity patterns. A loss of PE would cause a disproportionately large loss of evolutionary history (10, 11), making PE a key metric of biodiversity in conservation prioritization. However, global geographic patterns of PE and their past and present environmental drivers have yet to be explored in seed plants – a taxonomic group of fundamental ecological importance (12, 13).

Generally, endemism is shaped by multiple biogeographic and evolutionary processes (see Supplementary Table S1 for main hypotheses of PE determinants), and has been linked to limited gene flow and dispersal, which in turn foster speciation and limit range expansion. Dispersal limitation is promoted by geographic isolation due to physical and ecological dispersal barriers, e.g., oceans and mountain ranges or climatic gradients (14). Endemism on isolated islands exceeds that of mainland areas (2, 9), and also Mediterranean-type climate regions show high endemism due to local speciation events driven by their peculiar climate (15). Beyond isolation, long-term climatic stability coupled with habitat or resource heterogeneity may increase opportunities for speciation by allowing for the evolution of narrow physiological tolerances and ecological specialization (16, 17). In addition, endemism is facilitated by the longterm survival of spatially restricted species and their accumulation over long timescales. Consequently, climate, long-term climatic stability and environmental heterogeneity are hypothesized to affect the persistence of range-restricted species. Warm and humid climates may support larger populations in small regions by offering sufficient resources (18, 19). In addition, long-term climatically stable regions may serve as refugia for endemic species during periods of shifting climate (20, 21) and topographically heterogeneous regions require the species to move over only relatively short altitudinal distances in response to climate change reducing their extinction risk (22, 23).

Factors favoring the formation or persistence of endemic species do not need to be mutually exclusive. However, the influence of these processes may vary across the landscape over geological time, leading to assemblages of more recently evolved endemics or those that diverged longer ago, or both (24). During periods of pronounced climate change (e.g. Quaternary glacial cycles), plant distributions shifted greatly, resulting in repeated range contractions to suitable refugia followed by range expansions in more favorable periods (16, 25). After periods of climatically unfavorable conditions, not all plants could migrate out of their refugia to disperse to and fill their formerly occupied range (relictualization) (26, 27). Species that were once widespread and are now restricted to former refugia (“museums of biodiversity”) are called paleoendemics or relict endemics, which often represent old and range-restricted lineages (28, 29). In contrast, recently evolved lineages that are still restricted to their areas of origin like isolated oceanic islands or mountain regions (“cradles of biodiversity”) are called neoendemics (30, 31). Therefore, floras with high neo- or paleoendemism have likely been shaped by different processes affecting species diversification and persistence (see Supplementary Table S2 for main hypotheses of neo- and paleoendemism determinants).

Here, we reveal global patterns of PE for seed plants by integrating the globally most comprehensive regional plant inventory dataset (32, 33) across 912 geographical regions containing ~320,000 species (Supplementary Fig. S1) with a large phylogeny (34). We distinguish between centers of neoendemism and paleoendemism and centers of both types of endemism (11). Specifically, our aims are: (i) to reveal geographic patterns of PE for seed plants at the global scale; (ii) to test hypotheses related to isolation, climate, environmental heterogeneity and past climate change on global patterns of seed plant PE; (iii) to identify centers of neoendemism and paleoendemism across the world; (iv) and to assess how past environmental change and geological history affected the spatial concentration of neo- and paleoendemism.

## Results

### Global patterns and drivers of phylogenetic endemism

Phylogenetic endemism of seed plants varied greatly among regions and was highest on islands and in topographically heterogeneous tropical mainland regions (Fig. 1). To account for the bias introduced by the multitude of apomictic taxa in the temperate Northern Hemisphere, these and all other main results are based on the global flora of 273,838 seed plant species excluding all species from 293 genera that contain apomictic species (35) (see Methods for more details and supplementary material for results including apomictic species). Because PE depends on reliable range size estimates, we calculated PE based on two different ways to measure the range size of each species: (i) the total area (PE.area) of regions a species occurs in and (ii) the number of these regions (PE.count).

**Fig.1.**
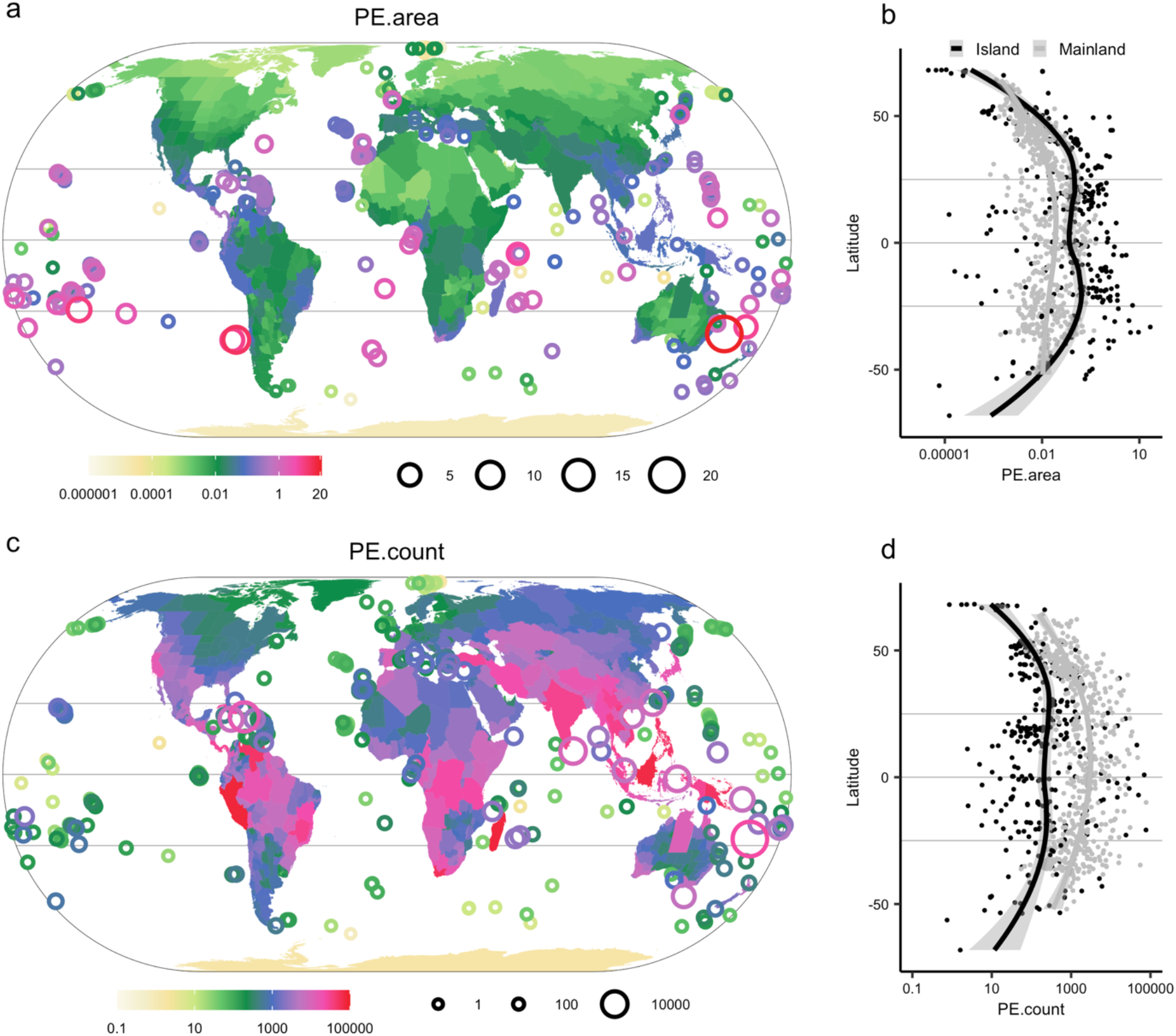
Global patterns of phylogenetic endemism of seed plants and its distribution along latitude. In a and b, phylogenetic endemism is calculated based on species range size measured as the area of regions where a species occurs (PE.area); In c and d, phylogenetic endemism is calculated based on species range size measured as the count of regions where a species occurs (PE.count). In b and d, the fitted lines are lowess regressions, separately fitted for islands and mainland regions. Log_10_ scale is used for phylogenetic endemism in all panels and maps use Eckert IV projection.

We found that PE.area was almost 15-fold higher on islands than in mainland regions (mean PE.area of islands and mainland regions: 0.44 vs 0.03 Myr·km^-2^). PE.area decreased from a mean value of 0.052 Myr·km^-2^ in mainland tropics to 0.018 Myr·km^-2^ in non-tropical mainland regions (Fig. 1a and b). Furthermore, PE.area peaked on subtropical islands located in the Southern Hemisphere, with Lord Howe Island having the highest PE.area overall (21.65 Myr·km^-2^), while Costa Rica showed the highest values among mainland regions (0.44 Myr·km^-2^; Table 1 and Supplementary Table S3). In contrast, PE.count peaked in the tropics both for islands (Madagascar: 76,964 Myr) and mainland regions (Peru: 77,729 Myr; Fig. 1c and d; Table 1).

**Table 1.** Top ten regions of phylogenetic endemism for seed plants. Phylogenetic endemism is calculated based on two different ways to measure range size of each species: (i) as the total area of regions a species occurs in (PE.area) and (ii) as the number of these regions (PE.count). See Supplementary Table S3 for top islands with different minimum of area size and Supplementary Table S4 for top ten regions based on the dataset including apomictic taxa.

| PE.area |  |  |  | PE.count |  |  |  |
| --- | --- | --- | --- | --- | --- | --- | --- |
| Mainland |  | Island |  | Mainland |  | Island |  |
| Costa Rica | 0.44 | Lord Howe Island | 21.65 | Peru | 77729 | Madagascar | 76964 |
| Pichincha, Ecuador | 0.44 | Masatierra | 9.37 | Western Cape Province, South Africa | 55197 | Borneo | 69591 |
| Western Cape Province, South Africa | 0.40 | Tubuai Island | 6.16 | Venezuela | 47547 | Papua New Guinea | 49859 |
| Panama | 0.40 | Masafuera | 5.34 | Vietnam | 34535 | Philippines | 45383 |
| Carchi, Ecuador | 0.30 | Norfolk Island | 4.78 | Thailand | 32192 | Indonesian New Guinea | 29119 |
| Valle del Cauca, Colombia | 0.24 | Silhouette | 3.92 | Minas Gerais | 30592 | Sumatra | 27187 |
| Antioquia, Colombia | 0.24 | Mahe | 2.92 | India | 30181 | New Caledonia | 24092 |
| Rio de Janeiro, Brazil | 0.23 | Henderson | 2.24 | Panama | 29108 | Cuba | 23675 |
| Imbabura, Ecuador | 0.22 | Fulanga | 2.09 | Yunnan | 28127 | Japan | 16511 |
| Cotopaxi, Ecuador | 0.22 | Rarotonga | 2.09 | Asiatic Turkey | 26746 | Hispaniola | 14738 |

The environmental factors we hypothesized to affect PE (i.e. isolation, climate, environmental heterogeneity and past climate change; Supplementary Table S1) explained 80.3% of the variance in PE.area and 88.1% in PE.count (Fig.2). The effects of environmental factors on PE were largely similar for the two ways of how range size was quantified (differing most prominently for region area which had a positive effect on PE.count and a negative effect on PE.area) (Fig. 2). Surrounding landmass proportion, a proxy for isolation, which is lowest for remote islands and highest for regions located in the centers of large continents (36), emerged as the primary driver of PE.area. As hypothesized, surrounding landmass proportion was negatively related to PE.area and PE.count (Fig.2a), indicating that high PE occurred on islands and in mainland regions that are partly surrounded by water bodies such as coastal regions or peninsulas. Also, as expected, PE strongly increased with elevational range and number of soil types (proxies of environmental heterogeneity). Among climatic factors, energy and water availability had strong associations with PE, with increasing length of growing season and mean annual temperature leading to higher PE. Climate seasonality (incl. temperature and precipitation seasonality), however, had no or only weak positive effects on PE. Relatively recent past climate change left prominent traces in PE, but this was not detectable for climate changes in deeper time. PE increased with temperature stability since the Last Glacial Maximum (LGM; 21 Ka), while velocity of temperature change since the LGM had a negative effect. However, we found no significant relationship between PE and temperature anomaly since the mid-Pliocene warm period (~3.264 – 3.025 Ma). To test if the effects of environmental predictors on PE varied between isolated regions (e.g. islands) and less isolated regions (e.g. mainland regions), we included interactions between each individual predictor and surrounding landmass proportion in the models. We found that the positive effect of mean annual temperature on PE increased with decreasing surrounding landmass proportion (Fig. 2b). Almost the same results of PE patterns and their drivers were found for the dataset including apomictic taxa (Supplementary Fig. S2, Fig. S3 and Table S4).

**Fig. 2.**
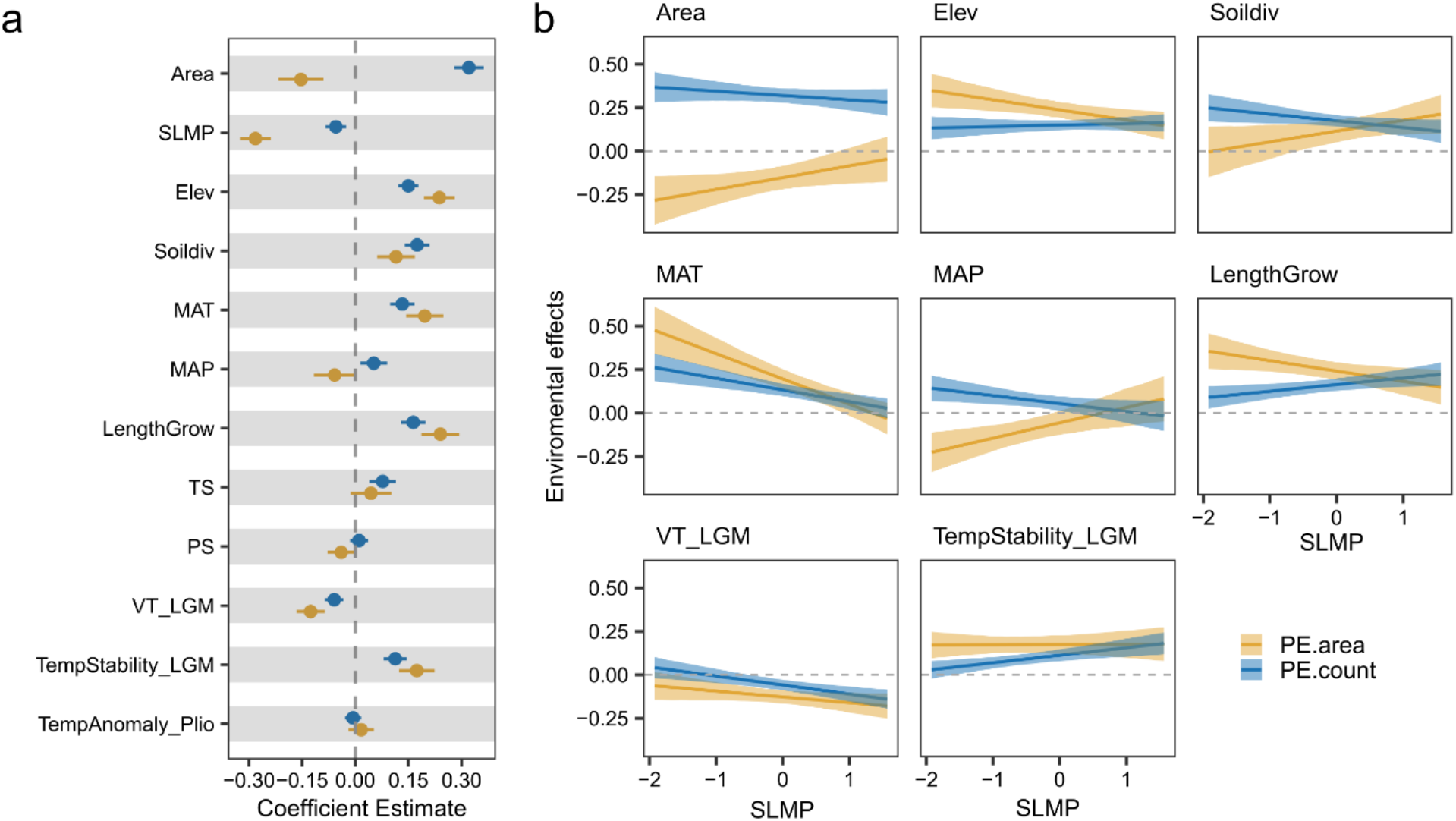
Determinants of phylogenetic endemism in seed plants based on spatial models including environmental factors and interactions between each environmental factor and surrounding landmass proportion. a, standardized regression coefficients of individual environmental factors. Bars around each point show the standard error of the coefficient estimate. b, significant interaction terms in the models visualized as effects of environmental factors on phylogenetic endemism (model coefficients on y-axis) with varying surrounding landmass proportion (x-axis). Lines and shadings represent 95% confidence intervals. Results are shown for phylogenetic endemism based on two competing ways of measuring range size of species. PE.area indicates phylogenetic endemism calculated based on range size of species as the area of regions where a species occurs, while PE.count is calculated based on range size of species as the count of these regions. Area = region area; SLMP = surrounding landmass proportion; Elev = elevational range; Soildiv = number of soil types; MAT = mean annual temperature; MAP= mean annual precipitation; LengthGrow = length of growing season; TS = temperature seasonality; PS = precipitation seasonality; VT_LGM = velocity of temperature change since the Last Glacial Maximum; TempStability_LGM = temperature stability since the Last Glacial Maximum; TempAnomaly_Plio = temperature anomaly between the mid-Pliocene warm period and present-day.

### Global centers and determinants of neo- and paleoendemism

We uncovered centers of evolutionarily old and range-restricted species, centers of evolutionarily young and range-restricted species as well as centers of both using a categorical analysis of neo- and paleoendemism (CANAPE) (11). We found that many remote islands (e.g. Mauritius and Juan Fernández Islands) and large continental fragment islands, such as Madagascar and Hispaniola, harbored both unusually high neo- and paleoendemism (i.e. centers of super-endemism) (Fig.3a and b). In contrast, New Caledonia as well as large continental islands in south-east Asia, such as New Guinea, Sumatra and Java, were identified as centers of paleoendemism. Mainland regions characterized by tropical moist forests, such as Amazonia, Peru, west Colombia, central Africa and Indochina, were found to harbor high numbers of paleoendemics. Mediterranean-climate regions stood out as extra-tropical hotspots of seed-plant PE among mainland regions. Regions in Central Chile and south-western Australia were characterized by both neo- and paleoendemic plants, while the Cape of south Africa, California and parts of the Mediterranean Basin (i.e. Spain and Asiatic Turkey) were centers of neoendemism. Some inconsistencies emerged for different measures of species range size. For example, the Himalaya was a center of neo- and paleoendemism based on PE.count, while it did not emerge as an endemism center based on PE.area. Comparing patterns including and excluding apomictic taxa, the most prominent differences occurred in European countries which were identified as endemism centers when including apomictic species, due to high numbers of apomictic range-restricted species in genera like *Rubus* and *Hieracium* (Supplementary Fig. S4 and S5).

**Fig. 3.**
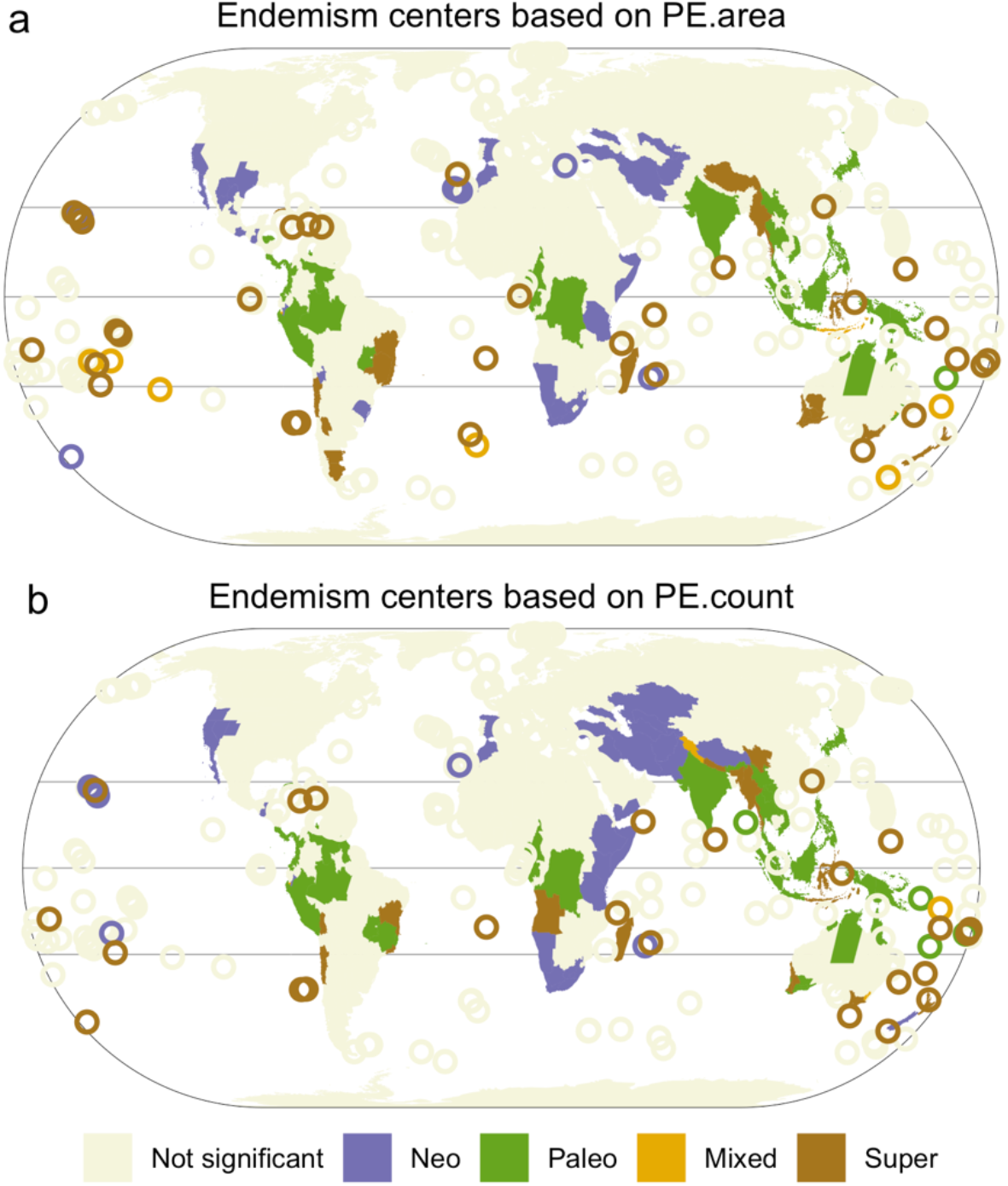
Global centers of neo- and paleoendemism for seed plants identified by the categorical analysis of neo- and paleoendemism (CANAPE). Colored regions present different types of endemism centers: violet, neoendemism; green, paleoendemism; yellow, mixed-endemism (i.e. neo- and paleoendemism); and brown indicating super-endemism (i.e. centers with both extremely high neo- and paleoendemism); beige, not significant. Patterns of neo- and paleoendemism are distinguished based on phylogenetic endemism with two competing ways of measuring species range size. PE.area indicates phylogenetic endemism calculated based on range size of species as the area of regions where a species occurs (a), while PE.count is calculated based on range size of species as the count of these regions (b). For patterns including apomictic taxa see Supplementary Fig. S4.

We assessed the impacts of geological history and past climate change on neo- and paleoendemism by modeling the standardized effect size of relative PE (see Methods for details) for regions which showed significantly high PE in response to past climatic and geological factors (Supplementary Table S2). We included the geographic type of each region (distinguishing between mainland regions and continental shelf islands, continental fragments and oceanic islands) and elevational range (to distinguish between mountainous and non-mountainous regions) to represent geological history. We found that oceanic islands showed significantly higher neoendemism than the mainland endemism centers identified in our study, while continental islands did not show a significant difference (Fig. 4). With increasing elevational range, neoendemism rose based on PE.count. However, this relationship was not significant based on PE.area. We found that centers of neo- and paleoendemism both had significantly higher elevational ranges than regions with low PE (Supplementary Fig. S6a and S7a). Past climate change was a major driver of neo-vs paleoendemism, with increasing temperature stability since the LGM increasing paleoendemism. Besides climate since the LGM, we also found significant relationships between temperature anomaly since the mid-Pliocene warm period and neo- and paleoendemism (Supplementary Fig. S6d and S7d). Specifically, super-endemic regions showed a significantly lower temperature anomaly since the mid-Pliocene than other regions. The relationships between the environmental variables and neo-vs paleoendemism were almost same for the dataset including apomictic taxa (Supplementary Fig. S8).

**Fig. 4.**
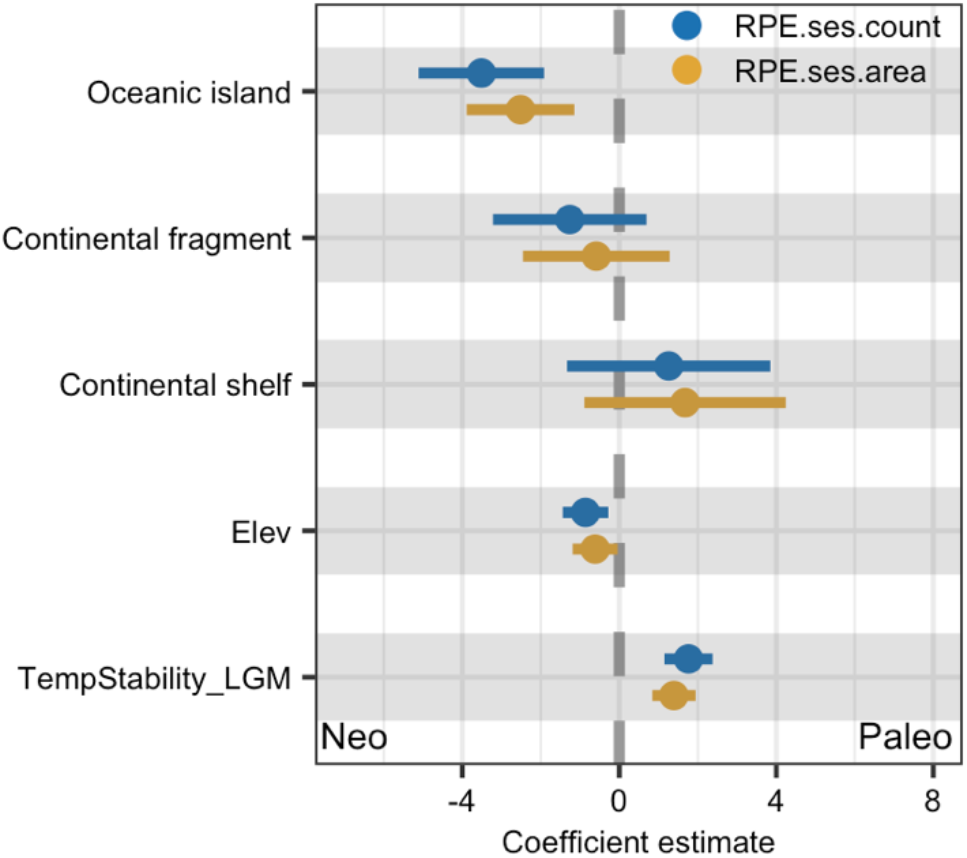
Determinants of neo- and paleoendemism. Standardized regression coefficients of environmental factors are shown from spatial models of the standardized effect size of relative phylogenetic endemism of seed plants for regions with significantly high phylogenetic endemism. A positive effect of environmental factors in the model represents higher paleoendemism along the environmental factor, while a negative effect represents higher neoendemism. The reference level of geographic type is mainland regions. RPE.ses.area indicates the standardized effect size of relative phylogenetic endemism calculated based on range size of species as the area of regions where a species occurs, while RPE.ses.count is calculated based on range size of species as the count of these regions.

## Discussion

Our study reveals islands and mountainous regions in the tropics and subtropics as global centers of phylogenetic endemism of seed plants. High numbers of geographically restricted and evolutionarily unique lineages, shaped by processes related to the generation and persistence of range-restricted species, led to varied concentrations of neo- and paleoendemism. We show that past and present environmental factors help to explain how diversification and relictualization (i.e. the persistence of species that went extinct elsewhere) shape the distribution of PE and neo- and paleoendemism worldwide. Understanding these drivers of PE and knowing particularly those regions with both high neo- and paleoendemism that act simultaneously as “museum” and “cradle” of biodiversity is of great importance for biodiversity conservation.

Geographic isolation resulted in high PE on islands, which may stem from *in-situ* speciation in isolation or relictualization (27). While speciation events require time for island species to evolve into phylogenetically distinct species, high PE may accumulate over shorter times through relictualization (37, 38), resulting from species extinctions on the mainland and other islands until those species only occur on few or even single islands. Relict lineages on islands may be old and only distantly related to other species on the same island while species that diversified on islands are often young and closely related. High PE on islands may hence be a result of a combination of diversification leading to neoendemism and relictualization leading to paleoendemism on islands. Additionally, relictualization may explain that ancient endemic species are sometimes even older than the formation of the island, such as the only member of the genus *Hillebrandia sandwicensis* on the Hawaiian Islands (39) and the only member of the oldest known angiosperm family (Amborellaceae) *Amborella trichopoda* on New Caledonia (40). Furthermore, diversification of island species is constrained by available resources and niches. For example, the probability of *in-situ* speciation scales positively with island size (41). This may explain the stronger effects of some environmental factors, such as energy availability and elevation range, on island PE than on mainland PE.

When comparing islands of different geological origin, we found that oceanic islands are characterized by higher neoendemism than continental islands, which may be explained by their unique geological history (27). Oceanic islands have not been connected with continental landmasses in the past but emerged from the oceans due to volcanic or tectonic activity. Untapped resources and the lack of enemies and competitors allowed plant species that colonized oceanic islands to diversify (14, 31). Considering the comparatively short geological lifespan of oceanic islands, speciation on these islands happened comparatively recently and species resulting from one lineage are still closely related, leading to neoendemism. However, some oceanic islands were identified as centers of super-endemism (e.g. Hawai’i Island), where relictualization and diversification happened in concert. Continental fragments and continental shelf islands, in contrast, were once parts of continents and are now separated from those by tectonic drift or sea-level rise. These islands were originally inhabited by floras comparable to those of the continents they were isolated from. The prolonged isolation (tens of million years) of continental fragments allowed for the accumulation of relict lineages as well as *in-situ* speciation, which led to high neo- and paleoendemism on some of these islands. The origins of endemism on large continental fragments are often still debated (42). Apart from more recent colonization events, evolution after vicariance or early long-distance dispersal events may have led to particularly old endemic species. For example, the majority of endemics on Madagascar evolved from lineages that originated from Cenozoic dispersal events (43), while few groups (e.g. the genus *Takhtajania*) date back to a potential Gondwanan vicariance (44). Also, islands located in south-east Asia showed high concentrations of paleoendemism, which is due to numerous relict lineages that have survived the last two mass extinctions (45). Consequently, our results reinforce the conservation urgency for islands which are often occupied by neoendemics as well as paleoendemics that represent millions of years of unique evolutionary history (46).

Tropical montane regions are well-known centers of taxonomic and phylogenetic plant diversity (e.g., ref. 47). Due to their complex topography and geologic and climatic histories, they also hold exceptionally narrow-ranged species (48). Consequently, montane regions, particularly tropical ones, emerged as centers of phylogenetic endemism for plants, as indicated by the positive effects of elevational range and diversity of soil types on PE. Importantly, we found that these regions accumulated both high neo- and paleoendemism. On the one hand, montane regions show remarkable diversification of their plant lineages and therefore foster high neoendemism, acting as “cradles” of biodiversity (49, 50). This diversification is the consequence of multiple mechanisms, including adaptation to diverse niches during long-term orogeny (50, 51), or divergence resulting from dynamic connectivity between habitats related to climatic fluctuation (52, 53). On the other hand, montane regions support the persistence of ancient lineages over time, acting as “museums” for paleoendemics (54). This results from steep environmental gradients with diverse microclimates in montane regions, allowing species to track their climate niche through altitudinal range shifts during climatic change periods (23).

Our results show how past climate has affected present patterns of PE, with climate stability since the Last Glacial Maximum promoting the accumulation of narrow-ranged species, especially for paleoendemics. Cooler temperatures during glacial periods may have caused range contractions and selective extinctions of narrow-ranged species, and thus likely removed or reduced their ranges in less stable regions (21). Additionally, recolonization of mostly wide-ranged species after glacial retreats may have further decreased PE in regions with historically variable climate (55). In contrast, some regions such as islands, coastal or montane regions have suffered less from past climatic changes because of the buffering effect of the oceans against climatic changes (28) and the topographically diverse microclimates in montane regions (23). Different from climate during the LGM, the mid-Pliocene warm period represented warmer climates compared to the current climate. Nevertheless, the high concentration of both neo- and paleoendemism in regions with less climatic changes since the mid-Pliocene warm period again emphasizes the vital roles of long-term climatic stability on speciation and persistence of narrow-ranged species. Beyond past climate, energy and water availability displayed a strong positive effect on PE of plants. This may be linked to lower extinction risks for narrow-ranged plants under warm and wet climates by offering favorable environments and sufficient resources for larger population sizes in smaller areas (56). Additionally, we found that Mediterranean-climate regions acted as extra-tropical hotspots of plant endemism, especially with high neoendemism. This may be attributed to the recent and rapid speciation in these regions, triggered by the unique climatic regime characterized by high seasonality and summer drought (15, 57).

Generally, larger regions host more endemics as well as wide-ranged species because of their overall higher plant diversity (47, 58). Here, we observed a negative association between region area and PE when species range sizes were measured as the total area of the occupied regions. Specifically, PE.area peaked on some small islands (e.g. Lord Howe Island) and showed lower values in large mainland regions. However, PE.area of large mainland regions was possibly underestimated because the range sizes of endemics that only occur in small suitable habitats within large regions were greatly overestimated. In contrast, there was a positive association between region area and PE when we measured species range sizes as the count of occupied regions. However, this method ignores the variation of area across regions and disregards that endemics of small regions likely have smaller ranges than endemics of larger regions leading to an underestimation of PE for small regions. Area therefore rather acted as a covariate to control for the underestimation or overestimation of PE in our two metrics than as an environmental predictor.

In conclusion, our study uncovers global patterns of phylogenetic endemism for seed plants and disentangles the complex joint contribution of isolation, heterogeneity, climate and long-term climatic stability. Integration of unprecedented phylogenetic information allowed us to distinguish global centers of neo- and paleoendemism, highlighting tropical mountains and oceanic and large continental islands as hotspots of evolutionarily distinct endemic species. These regions have experienced unique climatic and geological histories, which have driven important evolutionary and ecological processes of diversity generation and maintenance. Consequently, these regions are of crucial conservation value and need to be put under protection.

## Materials and Methods

### Species distribution data

We used regional species composition data for native seed plants from the Global Inventory of Floras and Traits (GIFT version 3.0: http://gift.uni-goettingen.de) (33) and the World Checklist of Vascular Plants (WCVP, http://wcvp.science.kew.org/) (32). GIFT contains regional plant inventories from published floras and checklists for ~ 3500 geographic regions worldwide representing islands, protected areas, biogeographical regions (e.g. botanical countries) and political units (e.g. countries, provinces). WCVP is a comprehensive taxonomic compilation of vascular plants and offers distribution information of species in WGSRPD Level-3 units (i.e. 369 botanical countries). We downloaded information for each non-synonym species in WCVP using the function *pow_lookup* in the R package *taxize* (59) and extracted their distribution and biogeographic status across all botanical countries. We then combined all native seed plant occurrences from WCVP with all native seed plant checklists from GIFT available for the same regions. To obtain finer-grain distribution information for some large regions, we replaced botanical countries by smaller regions from GIFT where available (e.g. the individual departments of Bolivia instead of the entire country). To replace large botanical countries by smaller regions from GIFT, we removed the larger regions only when smaller regions were nested within the larger regions and all nested regions completely covered the larger regions for mainland regions, and replaced archipelagos by individual islands if the individual islands made up most of the archipelago. Because all non-hybrid species names in GIFT 3.0 were standardized and validated based on taxonomic information provided by WCVP, we were able to directly combine WCVP and GIFT data. We retained taxonomically unmatched species names because of the low percentage of these species per region (i.e. 99.7% of all species names were taxonomically matched on average across regions). We excluded regions < 10 km^2^ (area not permanently covered by ice). All excluded regions were islands and only few excluded islands host endemic species (49 endemic species on 112 islands < 10 km^2^ in GIFT). The final dataset included 317,985 seed plant species for 912 geographic regions covering all landmass worldwide with varying area size from 10 to 3,069,766 km^2^ (median: 23,192 km^2^), consisting of 597 mainland regions and 315 islands or island groups (Supplementary Fig. S1).

### Apomictic taxa

Apomixis is a special case of uniparental reproduction via asexually formed seeds (60). Apomixis is tightly associated with hybridization and polyploidization, and may promote reticulate evolution and the formation of a multitude of novel lineages (60). European brambles (*Rubus* subgen. *Rubus*, Rosaceae), for example, consist mostly of apomictic taxa (only 4 out 748 accepted species are sexual) owing to speciation via reticulation and apomixis (61). However, taxonomic treatment of these complex groups of apomictic taxa and underlying species concepts are contentious. Additionally, regional floras and checklists differ in the level of detail at which these groups are included and taxonomically resolved. Consequently, the global distribution of apomictic taxa is geographically biased (particularly towards the well sampled floras of Europe), affecting the assessment of endemism, especially for regions with a high proportion of apomictic taxa. To account for the bias introduced by apomictic taxa, we repeated all analyses including and excluding all the species from 293 genera that contain apomictic species according to the Apomixis Database (http://www.apomixis.uni-goettingen.de) (35). The distribution dataset excluding apomictic species included 273,838 species and was used for the main analyses. The Apomixis Database has been constructed only for angiosperms. It has however been shown that apomixis is very rare in gymnosperms (60).

### Phylogeny

To measure phylogenetic endemism, we linked the species from the distribution dataset to a large, dated species-level phylogeny of seed plants with 353,185 tips (34). Not all species in the distribution dataset were included in the phylogeny. We therefore bound the remaining species to the congeners in the phylogeny by replacing all species of the genus with a polytomy, using the function *congeneric.merge* in the R package *pez* (62). We then excluded species not present in the distribution dataset from the phylogeny (hereafter called merged phylogeny). The merged phylogeny included 267,105 out of 273,838 species (97.5 %) in the dataset. Adding phylogenetic uncertainties via polytomies is a common issue when working with large phylogenetic trees (47, 63). However, phylogenetic metrics based on phylogenetic trees with higher numbers of polytomies have been shown to be highly correlated with metrics based on trees without or with fewer polytomies (64). To test whether adding species and replacing their genera by polytomies affects PE patterns, we built an additional phylogeny for sensitivity analyses. Specifically, all species from the original phylogeny that did not exist in the distribution dataset were excluded from the original phylogeny (hereafter called matched phylogeny). The matched phylogeny contained 212,525 species (77.6 % of all species in the distribution dataset). PE derived from the merged phylogeny was highly correlated to PE based on the matched phylogeny (Pearson’s r = 0.98), suggesting that calculating PE based on the merged phylogeny was robust. Identical procedures were also carried out for the datasets including apomictic species (Pearson’s r = 0.99). We used the merged phylogeny for all further analyses.

### Phylogenetic endemism

To investigate the distribution of seed plant endemism worldwide, we calculated phylogenetic endemism for each region following Rosauer et al. (10), as the sum of branch lengths of all species coexisting in a region, based on a phylogeny where each branch length is divided by the global range size of the species that descended from the branch. Because PE depends on reliable range size estimates, we measured the range size of each species and of each branch in two different ways: (i) as the number of regions a species occurs in (PE.count) and (ii) as the total area (not permanently covered by ice) of these regions (PE.area). PE.count overestimates PE particularly for large regions, since it disregards that the ranges of species endemic to small regions are likely smaller than the ranges of species endemic to larger regions. In contrast, PE.area accounts for the varied areas of regions in our dataset, but likely underestimates PE for large regions because their areas may be larger than the actual ranges of the species occurring inside. Despite the potential biases of both methods, the actual ranges and hence endemism fall within the range that is estimated by the two methods (Supplementary Fig. S9; see discussion for more details). We therefore repeated all analyses based on both metrics, considered those results particularly robust that emerged for both metrics and discussed differences critically.

### Neo- and paleoendemism

We used the categorical analysis of neo- and paleoendemism (CANAPE) (11) to distinguish between centers of neoendemism and paleoendemism. CANAPE is based on the assessment of statistical significance of PE and relative phylogenetic endemism (RPE). RPE is the ratio of PE measured on the actual phylogenetic tree divided by PE measured on a comparison tree that retains the actual tree topology but with all branches having the same length (11). Therefore, RPE allowed us to examine the degree to which branch lengths and hence clade ages matter for the observed patterns of PE. We carried out the CANAPE analysis for PE.count and PE.area, respectively. To test for significance of the metrics, we ran 1000 null model randomizations. In the null models, species occurrences across regions were randomly reassigned without replacement, keeping the species number in each region and the total number of regions occupied by each species constant (11). Distributions of null model values for each region were then used for non-parametric tests for significance of the observed values of the tested metric and also for calculating the standardized effect size of RPE. If the observed value of the tested metric fell into the highest 2.5% or lowest 2.5% of the null distribution for a region, it was identified as statistically significantly high or low, respectively (two-tailed test, α=0.05). This randomization-based significance test was carried out for PE measured on the actual tree (numerator of RPE), PE measured on the comparison tree with equal branch lengths (denominator of RPE), and RPE.

We then followed a two-step process to distinguish different centers of endemism following Mishler et al. (11). First, we identified regions with significantly high PE by testing whether PE measured on the actual tree (numerator of RPE), PE measured on the comparison tree with equal branch length (denominator of RPE), or both were significantly higher than expected (observed value > 95% of the randomization values; one-tailed test, α= 0.05). Second, we divided regions with significantly high PE into four categories of centers of endemism (paleo-, neo-, mixed, and super-endemism). If RPE of a region was significantly high or low (two-tailed test, α= 0.05), the region was defined as a center of paleoendemism or neoendemism, respectively. If the RPE was not significantly high or low, but both numerator and denominator of RPE were significantly high (α= 0.05), the region was defined as a center of mixed endemism. If a mixed endemism region had both a significantly high numerator and denominator of RPE at the α= 0.01 level, the region was identified as a center of super-endemism.

We also calculated the standardized effect size of relative phylogenetic endemism (RPE.ses) based on the null distributions of RPE obtained from the null model. RPE.ses was calculated as the difference between the observed values and the mean of the null distribution divided by the standard deviation of the null distribution. In contrast to the non-parametric test in CANAPE, RPE.ses quantifies, for each region, the degree to which disproportionately young or old lineages (i.e. shorter or longer branches) are spatially restricted. When only considering regions with significantly high PE, lower values of RPE.ses represent more young or younger lineages that are spatially restricted, while higher values represent more old or older lineages than expected by chance. Thus, this metric offers an opportunity to model and explore the relationship between the historical drivers and the spatial concentration of neo-vs. paleoendemism as a continuous variable. However, it should be noted that a region with both high neo- and paleoendemism may show a value close to zero.

### Predictors of phylogenetic endemism

We hypothesized that phylogenetic endemism is shaped by biogeographic and evolutionary processes that promote the origin and maintenance of range-restricted lineages. We identified a set of candidate predictor variables representing these processes and classified them into five categories: isolation, environmental heterogeneity, energy and water availability, climatic seasonality and long-term climatic stability (Supplementary Table S1). These factors have been shown or hypothesized to contribute to geographic patterns of plant endemism in previous studies (2, 13, 21, 54). We measured geographic isolation as the sum of the proportions of landmass area in the surrounding of the target regions within buffer distances of 100 km, 1000 km, and 10,000 km (36). Its value is lowest for remote islands and highest for regions located in the centers of large continents. We considered the number of soil types (65) and elevational range (66) for each region as proxies for environmental heterogeneity. We also included five ecologically relevant climatic variables representing main aspects of climate hypothesized to be important for plant endemism, namely mean annual temperature, mean annual precipitation, length of the growing season (i.e. number of days with temperatures exceeding a threshold of 0.9°C, without snow cover, and with sufficient soil water), temperature seasonality (i.e. standard deviation of mean monthly temperature × 100) and precipitation seasonality (i.e. coefficient of variation in monthly precipitation) (67). Climatic variables were extracted as mean values per region from the input raster layers.

To determine the contribution of long-term climatic stability to PE, we calculated temperature stability since the LGM (21 Ka), velocity of temperature change since the LGM, and temperature anomaly since the mid-Pliocene warm period (~3.264 – 3.025 Ma). The LGM and the mid-Pliocene warm period represent cooler and warmer climates compared to the current climates, respectively. Temperature stability since the LGM was calculated using the *climateStablity* R package (68). It calculates temperature differences between 1000 year time slices expressed as standard deviation and averages the results across all time slices. Temperature stability is then calculated as the inverse of the mean standard deviation rescaled to [0,1] (68). In addition, we calculated the velocity of temperature change since the LGM as the ratio between temporal change and contemporary spatial change in temperature, representing the speed with which a species would have to move its range to track analogous climatic conditions (54). Temperature anomaly since the mid-Pliocene warm period was calculated as the absolute difference in mean annual temperature between the mid-Pliocene warm period and present-day (69).

### Predictors of neo- and paleoendemism

Neo- and paleoendemism are hypothesized to be driven primarily by the geological history of a region and by past climate change or stability (Supplementary Table S2). We therefore included the geographic type of each region (distinguishing between mainland regions and continental shelf islands, continental fragments and oceanic islands) instead of surrounding landmass proportion, and elevational range (to distinguish between mountainous and non-mountainous regions) to represent geological history (33). We removed three islands with heterogeneous geologic origin from further analyses on neo- and paleoendemism. To test for the impacts of past climate change on neo- and paleoendemism, we included the variables of long-term climatic stability introduced above.

### Models of phylogenetic endemism

To assess the relationships between PE and environmental predictor variables, we fitted linear models with PE as a response variable. Beyond all predictor variables hypothesized to be important to PE (Supplementary Table S1), we included area size (km^2^) to control for the over- and underestimation of PE in large regions for PE.count and PE.area, respectively. We excluded regions with incomplete data of the predictor variables, leading to a dataset including 818 regions (incl. 236 islands and 582 mainland regions; see Supplementary Fig. S10 for correlations between predictors). PE was Log_10_-transformed prior to modeling. Some predictor variables (i.e. region area, elevational range, number of soil types, mean annual precipitation, temperature seasonality, precipitation seasonality, velocity in temperature change since the LGM, temperature stability since the LGM and temperature anomaly since the mid-Pliocene warm period) were also Log_10_-transformed to reduce skewness of their distributions. All continuous predictor variables were then standardized to zero mean and unit variance to aid model fitting and make their parameter estimates comparable. To test whether the effects of environmental predictors on PE differ for isolated islands compared to less isolated mainland regions, we included the interaction between each individual predictor and surrounding landmass proportion. To test if including surrounding landmass proportion correctly encapsulated the effect of insularity, we updated the model by replacing surrounding landmass proportion with a categorical variable indicating whether or not a region is an island. Since these models performed worse than models including surrounding landmass proportion (Supplementary Table S5), we retained surrounding landmass proportion for all further analyses.

Species distributions, environmental predictors and model residuals are often spatially autocorrelated, which may lead to biased parameter estimates and the violation of statistical assumptions (70). As spatial autocorrelation was detected in the model residuals (Supplementary Fig. S11a, b), we included a spatial autocovariate that represents the spatial autocorrelation in the residuals of non-spatial models (residual autocovariate models, RAC) (71). This autocovariate term was implemented as a spatial weight matrix of non-spatial model residuals based on an optimized neighborhood structure. Because most of our regions are political units with varying geometry and size, we used a sphere of influence to identify neighbors for each region (72). The sphere of influence for each focal region was defined as a circle around the centroid of a focal region within a radius equal to the distance to the centroid of the nearest neighboring region. When the sphere of influence of two regions overlapped, the two regions were considered neighbors. Overall, the RAC models successfully removed spatial autocorrelation from model residuals (Supplementary Fig. S11c).

### Statistical analyses of neo- and paleoendemism

To explore the potential drivers of spatial concentrations of neo- and paleoendemism, we fitted ordinary linear models to explain the variation in RPE.ses only for regions which showed significantly high PE based on CANAPE (Fig. 3 and Supplementary Fig. S4). We removed velocity of temperature change since the LGM and temperature anomaly since the mid-Pliocene warm period because of their low explanatory power for RPE.ses based on AIC values. Predictors retained for modeling contained all three aspects (i.e. islands, mountains and past climate change) hypothesized to affect neo- and paleoendemism. Likewise, we fitted spatial models by including a spatial autocovariate to remove spatial autocorrelation present in the residuals of the non-spatial models (Supplementary Fig. S12).

In addition, we compared the distribution of environmental factors for all regions (912 regions) across all CANAPE categories (i.e. neo, paleo, mixed, super-endemic and non-significant). Because the environmental factors were not normally distributed for each category separately, we used a non-parametric Kruskal-Wallis test followed by Wilcoxon pairwise comparisons (two-tailed tests with Holm’s correction) to identify which categories were different from each other (48). We repeated all modeling procedures for two PE metrics (PE.area and PE.count) and the dataset excluding and including apomictic taxa.

## Acknowledgements

L.C. was supported by China Scholarship Council (CSC) Grant (No.201808330443). P.P. and J.P. were supported by EXPRO grant no. 19-28807X (Czech Science Foundation) and long-term research development project RVO 67985939 (Czech Academy of Sciences). M.W. acknowledges DFG funding via iDiv (DFG FZT 118, 202548816). M.v.K. acknowledges DFG funding (grant 264740629). A.T. is also supported by DFG funding (grant 447332176). F.E. appreciates funding by Austrian Science Foundation FWF (Global Plant Invasions, grant no. I 5825-B). This work used the Scientific Compute Cluster at GWDG, the joint data center of Max Planck Society for the Advancement of Science (MPG) and University of Göttingen.

## Author contributions

L.C., H.K. and P.W. conceived the idea and developed the conceptual framework of the study. L.C., H.K., A.T., J.S., W.D., F.E., M.v.K., J.P., P.P., M.W. and P.W. were involved in collecting the data. L.C. performed the statistical analyses and wrote the first draft of the manuscript. L.C., H.K., A.T., J.S., W.D., F.E., M.v.K., J.P., P.P., M.W. and P.W. contributed to the writing and interpretation of the results.

## Reference

1. S. N. Sheth, N. Morueta-Holme, A. L. Angert, Determinants of geographic range size in plants. New Phytol. 226, 650–665 (2020).

2. G. Kier, et al., A global assessment of endemism and species richness across island and mainland regions. Proc. Natl. Acad. Sci. 106, 9322–9327 (2009).

3. N. Myers, R. A. Mittermeier, C. G. Mittermeier, G. A. B. da Fonseca, J. Kent, Biodiversity hotspots for conservation priorities. Nature 403, 853–858 (2000).

4. N. C. A. Pitman, P. M. Jørgensen, Estimating the size of the world’s threatened flora. Science 298, 989–989 (2002).

5. N. J. B. Isaac, S. T. Turvey, B. Collen, C. Waterman, J. E. M. Baillie, Mammals on the EDGE: conservation priorities based on threat and phylogeny. PLoS ONE 2, e296 (2007).

6. G. M. Mace, J. L. Gittleman, A. Purvis, Preserving the tree of life. Science 300, 1707–1709 (2003).

7. A. Purvis, P.-M. Agapow, J. L. Gittleman, G. M. Mace, Nonrandom extinction and the loss of evolutionary history. Science 288, 328–330 (2000).

8. D. P. Faith, Conservation evaluation and phylogenetic diversity. Biol. Conserv. 61, 1–10 (1992).

9. S. Veron, et al., High evolutionary and functional distinctiveness of endemic monocots in world islands. Biodivers. Conserv. 30, 3697–3715 (2021).

10. D. Rosauer, S. W. Laffan, M. D. Crisp, S. C. Donnellan, L. G. Cook, Phylogenetic endemism: a new approach for identifying geographical concentrations of evolutionary history. Mol. Ecol. 18, 4061–4072 (2009).

11. B. D. Mishler, et al., Phylogenetic measures of biodiversity and neo- and paleo-endemism in Australian Acacia. Nat. Commun. 5, 4473 (2014).

12. G. Murali, R. Gumbs, S. Meiri, U. Roll, Global determinants and conservation of evolutionary and geographic rarity in land vertebrates. Sci. Adv. 7, eabe5582 (2021).

13. B. Sandel, et al., Current climate, isolation and history drive global patterns of tree phylogenetic endemism. Glob. Ecol. Biogeogr. 29, 4–15 (2020).

14. M. Fernández-Mazuecos, et al., The radiation of Darwin’s giant daisies in the Galápagos Islands. Curr. Biol. 30, 4989–4998 (2020).

15. J. D. Thompson, S. Lavergne, L. Affre, M. Gaudeul, M. Debussche, Ecological differentiation of Mediterranean endemic plants. Taxon 54, 967–976 (2005).

16. M. Dynesius, R. Jansson, Evolutionary consequences of changes in species’ geographical distributions driven by Milankovitch climate oscillations. Proc. Natl. Acad. Sci. 97, 9115–9120 (2000).

17. R. Jansson, M. Dynesius, The fate of clades in a world of recurrent climatic change: Milankovitch oscillations and evolution. Annu. Rev. Ecol. Syst. 33, 741–777 (2002).

18. W. Jetz, C. Rahbek, R. K. Colwell, The coincidence of rarity and richness and the potential signature of history in centres of endemism. Ecol. Lett. 7, 1180–1191 (2004).

19. D. F. Rosauer, W. Jetz, Phylogenetic endemism in terrestrial mammals. Glob. Ecol. Biogeogr. 24, 168–179 (2015).

20. B. J. Enquist, et al., The commonness of rarity: Global and future distribution of rarity across land plants. Sci. Adv. 5, eaaz0414 (2019).

21. R. Jansson, Global patterns in endemism explained by past climatic change. Proc. R. Soc. Lond. B Biol. Sci. 270, 583–590 (2003).

22. K. D. Bennett, P. C. Tzedakis, K. J. Willis, Quaternary refugia of north European trees. J. Biogeogr. 18, 103–115 (1991).

23. A. S. Jump, C. Mátyás, J. Peñuelas, The altitude-for-latitude disparity in the range retractions of woody species. Trends Ecol. Evol. 24, 694–701 (2009).

24. L.-M. Lu, et al., Evolutionary history of the angiosperm flora of China. Nature 554, 234–238 (2018).

25. M. B. Davis, R. G. Shaw, Range shifts and adaptive responses to Quaternary climate change. Science 292, 673–679 (2001).

26. M. D. Crisp, S. Laffan, H. P. Linder, A. Monro, Endemism in the Australian flora. J. Biogeogr. 28, 183–198 (2001).

27. R. G. Gillespie, G. K. Roderick, Arthropods on islands: colonization, speciation, and conservation. Annu. Rev. Entomol. 47, 595–632 (2002).

28. Q. C. B. Cronk, Islands: stability, diversity, conservation. Biodivers. Conserv. 6, 477–493 (1997).

29. J. Fjeldsa°, J. C. Lovett, Geographical patterns of old and young species in African forest biota: the significance of specific montane areas as evolutionary centres. Biodivers. Conserv. 6, 325–346 (1997).

30. C. E. Hughes, G. W. Atchison, The ubiquity of alpine plant radiations: from the Andes to the Hengduan Mountains. New Phytol. 207, 275–282 (2015).

31. J. B. Losos, R. E. Ricklefs, Adaptation and diversification on islands. Nature 457, 830–836 (2009).

32. R. Govaerts, E. Nic Lughadha, N. Black, R. Turner, A. Paton, The World Checklist of Vascular Plants, a continuously updated resource for exploring global plant diversity. Sci. Data 8, 215 (2021).

33. P. Weigelt, C. König, H. Kreft, GIFT – A global inventory of floras and traits for macroecology and biogeography. J. Biogeogr. 47, 16–43 (2020).

34. S. A. Smith, J. W. Brown, Constructing a broadly inclusive seed plant phylogeny. Am. J. Bot. 105, 302–314 (2018).

35. D. Hojsgaard, S. Klatt, R. Baier, J. G. Carman, E. Hörandl, Taxonomy and biogeography of apomixis in angiosperms and associated biodiversity characteristics. Crit. Rev. Plant Sci. 33, 414–427 (2014).

36. P. Weigelt, H. Kreft, Quantifying island isolation–insights from global patterns of insular plant species richness. Ecography 36, 417–429 (2013).

37. Q. C. B. Cronk, Relict floras of Atlantic islands: patterns assessed. Biol. J. Linn. Soc. 46, 91–103 (1992).

38. Q. C. B. Cronk, The history of endemic flora of St Helena: a relictual series. New Phytol. 105, 509–520 (1987).

39. W. L. Clement, et al., Phylogenetic position and biogeography of *Hillebrandia sandwicensis* (Begoniaceae): a rare Hawaiian relict. Am. J. Bot. 91, 905–917 (2004).

40. L. Zhang, et al., The water lily genome and the early evolution of flowering plants. Nature 577, 79–84 (2020).

41. Y. Kisel, T. G. Barraclough, Speciation has a spatial scale that depends on levels of gene flow. Am. Nat. 175, 316–334 (2010).

42. A. Antonelli, et al., Madagascar’s extraordinary biodiversity: Evolution, distribution, and use. Science 378, eabf0869 (2022).

43. A. D. Yoder, M. D. Nowak, Has vicariance or dispersal been the predominant biogeographic force in Madagascar? Only time will tell. Annu. Rev. Ecol. Evol. Syst. 37, 405–431 (2006).

44. S. Buerki, D. S. Devey, M. W. Callmander, P. B. Phillipson, F. Forest, Spatio-temporal history of the endemic genera of Madagascar. Bot. J. Linn. Soc. 171, 304–329 (2013).

45. Ş. Procheş, S. Ramdhani, S. J. Perera, J. R. Ali, S. Gairola, Global hotspots in the present-day distribution of ancient animal and plant lineages. Sci. Rep. 5, 15457 (2015).

46. J. M. Fernández-Palacios, et al., Scientists’ warning – The outstanding biodiversity of islands is in peril. Glob. Ecol. Conserv. 31, e01847 (2021).

47. L. Cai, et al., Global models and predictions of plant diversity based on advanced machine learning techniques. New Phytol., nph.18533 (2022).

48. L. M. J. Dagallier, et al., Cradles and museums of generic plant diversity across tropical Africa. New Phytol. 225, 2196–2213 (2020).

49. V. S. F. T. Merckx, et al., Evolution of endemism on a young tropical mountain. Nature 524, 347–350 (2015).

50. Y. Xing, R. H. Ree, Uplift-driven diversification in the Hengduan Mountains, a temperate biodiversity hotspot. Proc. Natl. Acad. Sci. 114, E3444–E3451 (2017).

51. A. Antonelli, et al., Geological and climatic influences on mountain biodiversity. Nat. Geosci. 11, 718–725 (2018).

52. S. G. A. Flantua, et al., Snapshot isolation and isolation history challenge the analogy between mountains and islands used to understand endemism. Glob. Ecol. Biogeogr. 29, 1651–1673 (2020).

53. C. Rahbek, et al., Building mountain biodiversity: Geological and evolutionary processes. Science 365, 1114–1119 (2019).

54. B. Sandel, et al., The influence of Late Quaternary climate-change velocity on species endemism. Science 334, 660–664 (2011).

55. T. J. Davies, A. Purvis, J. L. Gittleman, Quaternary climate change and the geographic ranges of mammals. Am. Nat. 174, 297–307 (2009).

56. K. L. Evans, P. H. Warren, K. J. Gaston, Species–energy relationships at the macroecological scale: a review of the mechanisms. Biol. Rev. 80, 1–25 (2005).

57. L. M. Valente, V. Savolainen, P. Vargas, Unparalleled rates of species diversification in Europe. Proc. R. Soc. B Biol. Sci. 277, 1489–1496 (2010).

58. D. Storch, P. Keil, W. Jetz, Universal species–area and endemics–area relationships at continental scales. Nature 488, 78–81 (2012).

59. S. A. Chamberlain, E. Szöcs, taxize: taxonomic search and retrieval in R. F1000Research 2, 191 (2013).

60. L. Majeský, F. Krahulec, R. J. Vašut, How apomictic taxa are treated in current taxonomy: A review. Taxon 66, 1017–1040 (2017).

61. M. Sochor, R. J. Vašut, T. F. Sharbel, B. Trávníček, How just a few makes a lot: Speciation via reticulation and apomixis on example of European brambles *(Rubus* subgen. Rubus, Rosaceae). Mol. Phylogenet. Evol. 89, 13–27 (2015).

62. W. D. Pearse, et al., pez : phylogenetics for the environmental sciences. Bioinformatics 31, 2888–2890 (2015).

63. H. Qian, R. E. Ricklefs, W. Thuiller, Evolutionary assembly of flowering plants into sky islands. Nat. Ecol. Evol. 5, 640–646 (2021).

64. H. Qian, Y. Jin, Are phylogenies resolved at the genus level appropriate for studies on phylogenetic structure of species assemblages? Plant Divers. 43, 255–263 (2021).

65. T. Hengl, et al., SoilGrids250m: Global gridded soil information based on machine learning. PLoS ONE 12, e0169748 (2017).

66. J. J. Danielson, D. B. Gesch, “Global multi-resolution terrain elevation data 2010 (GMTED2010)” (2011) https://doi.org/10.3133/ofr20111073.

67. D. N. Karger, et al., Climatologies at high resolution for the earth’s land surface areas. Sci. Data 4, 170122 (2017).

68. H. L. Owens, R. Guralnick, climateStability: an R package to estimate climate stability from time-slice climatologies. Biodivers. Inform. 14, 8–13 (2019).

69. J. L. Brown, D. J. Hill, A. M. Dolan, A. C. Carnaval, A. M. Haywood, PaleoClim, high spatial resolution paleoclimate surfaces for global land areas. Sci. Data 5, 1–9 (2018).

70. C. Dormann, et al., Methods to account for spatial autocorrelation in the analysis of species distributional data: a review. Ecography 30, 609–628 (2007).

71. B. Crase, A. C. Liedloff, B. A. Wintle, A new method for dealing with residual spatial autocorrelation in species distribution models. Ecography 35, 879–888 (2012).

72. J. Y. Lim, J.-C. Svenning, B. Göldel, S. Faurby, W. D. Kissling, Frugivore-fruit size relationships between palms and mammals reveal past and future defaunation impacts. Nat. Commun. 11, 4904 (2020).

