## Supplementary material for "Phylogenetic endemism of the world’s seed plants"

### **Supplementary material for**
Phylogenetic endemism of the world's seed plants.

Lirong Cai*, Holger Kreft, Amanda Taylor, Julian Schrader, Wayne Dawson, Franz Essl, Mark van Kleunen, Jan Pergl, Petr Pyšek, Marten Winter, Patrick Weigelt

*Lirong Cai


**This PDF file includes:**

Figures S1 to S12
Tables S1 to S5
References S1
SI References

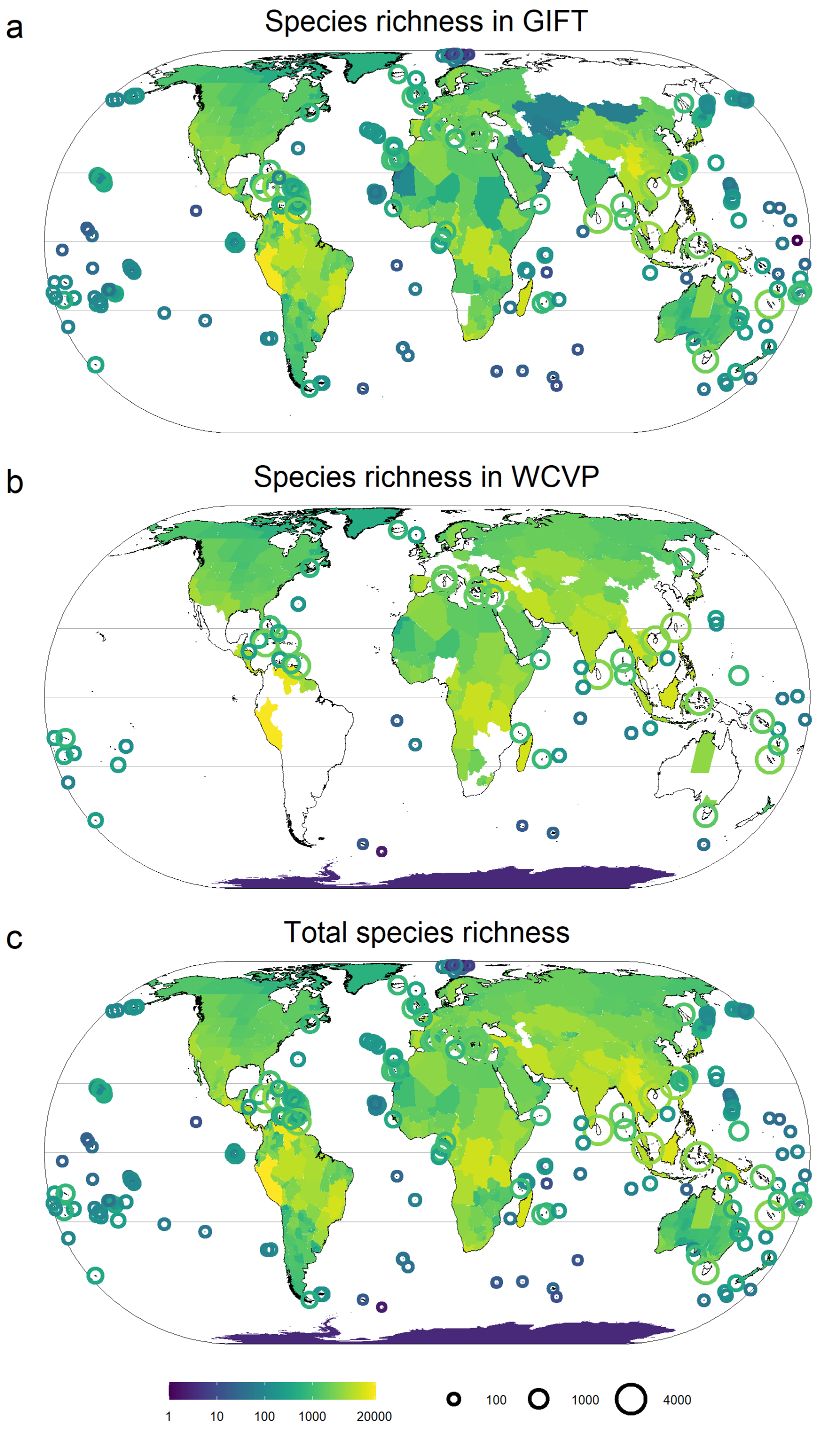


**Fig. S1** Observed species richness of seed plants for 912 geographic regions from (a) the Global Inventory of Floras and Traits (GIFT) (1) and (b) the World Checklist of Vascular Plants (WCVP) (2), combined in (c) to estimate phylogenetic endemism at the global scale and as fine grain as possible. $Log_{10}$ scale is used for species richness and maps use Eckert IV projection.

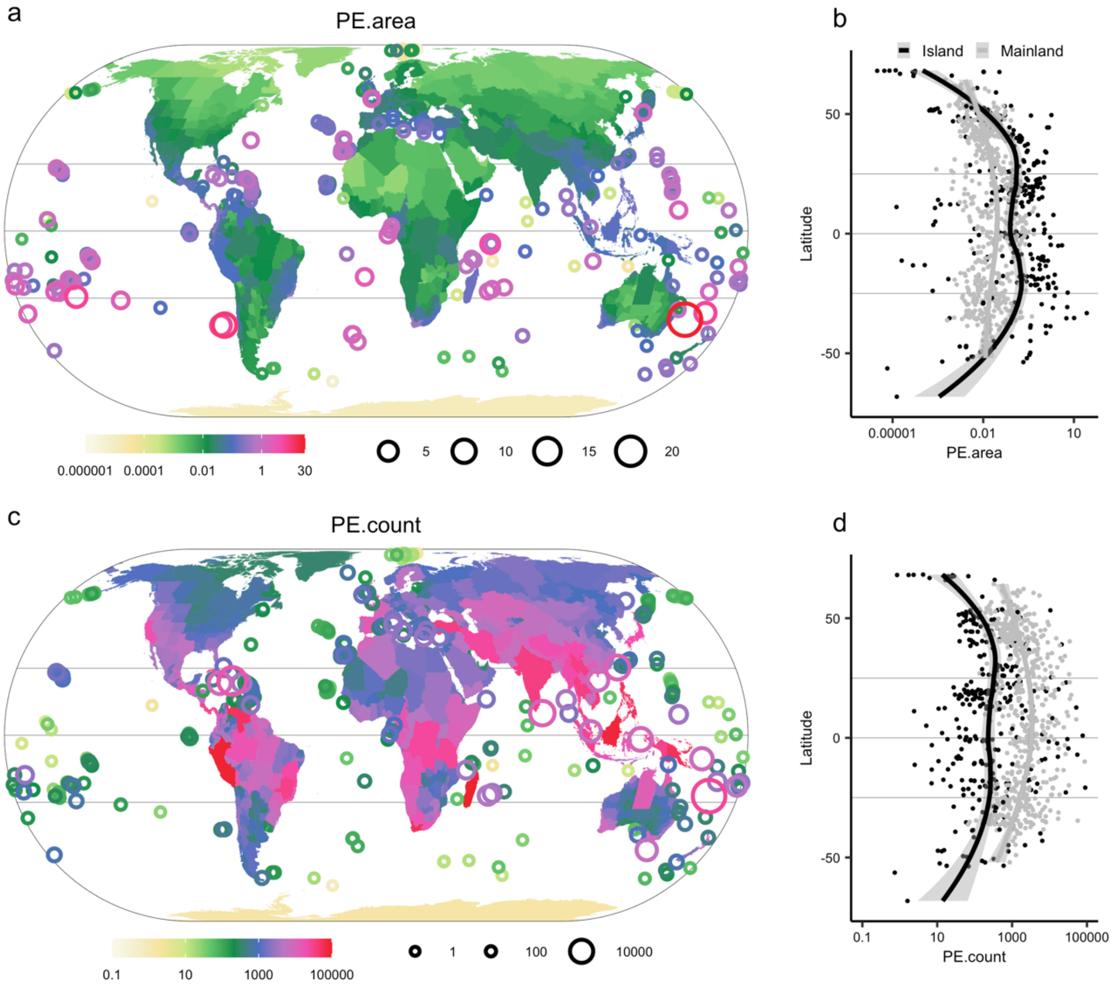


**Fig. S2** Global patterns of phylogenetic endemism of seed plants and its distribution along latitude based on the distribution data including apomictic taxa. In a and b, phylogenetic endemism is calculated based on species range size measured as the area of regions where a species occurs (PE.area); In c and d, phylogenetic endemism is calculated based on species range size measured as the count of regions where a species occurs (PE.count). In b and d, the fitted lines are lowess regressions, separately fitted for islands and mainland regions. $Log_{10}$ scale is used for phylogenetic endemism in all panels and maps use Eckert IV projection.

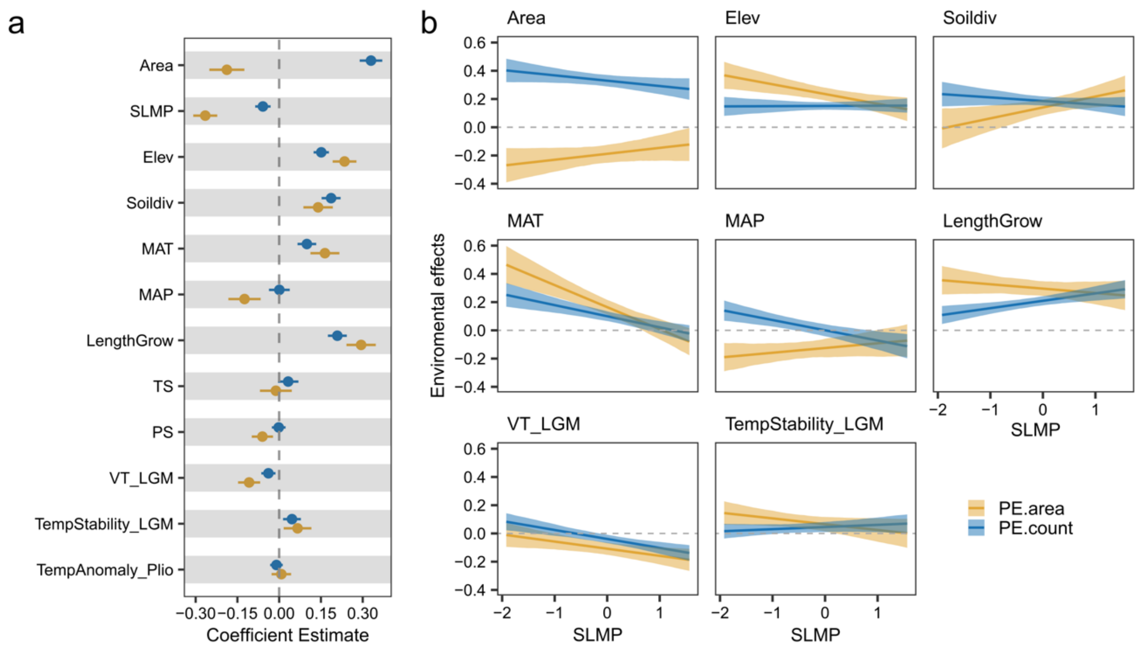


**Fig. S3** Determinants of phylogenetic endemism in seed plants based on the distribution data including apomictic taxa. Results are obtained from spatial models including environmental factors and interactions between each environmental factor and surrounding landmass proportion. a, standardized regression coefficients of individual environmental factors. Bars around each point show the standard error of the coefficient estimate. b, significant interaction terms in the models visualized as effects of environmental factors on phylogenetic endemism (model coefficients on y-axis) with varying surrounding landmass proportion (x-axis). Lines and shadings represent 95% confidence intervals. Results are shown for phylogenetic endemism based on two competing ways of measuring range size of species. PE.area indicates phylogenetic endemism calculated based on range size of species as the area of regions where a species occurs, while PE.count is calculated based on range size of species as the count of these regions. Area = region area; SLMP = surrounding landmass proportion; Elev = elevational range; Soildiv = number of soil types; MAT = mean annual temperature; MAP= mean annual precipitation; LengthGrow = length of growing season; TS = temperature seasonality; PS = precipitation seasonality; VT_LGM = velocity of temperature change since the Last Glacial Maximum; TempStability_LGM = temperature stability since the Last Glacial Maximum; TempAnomaly_Plio = temperature anomaly between the mid-Pliocene warm period and present-day.

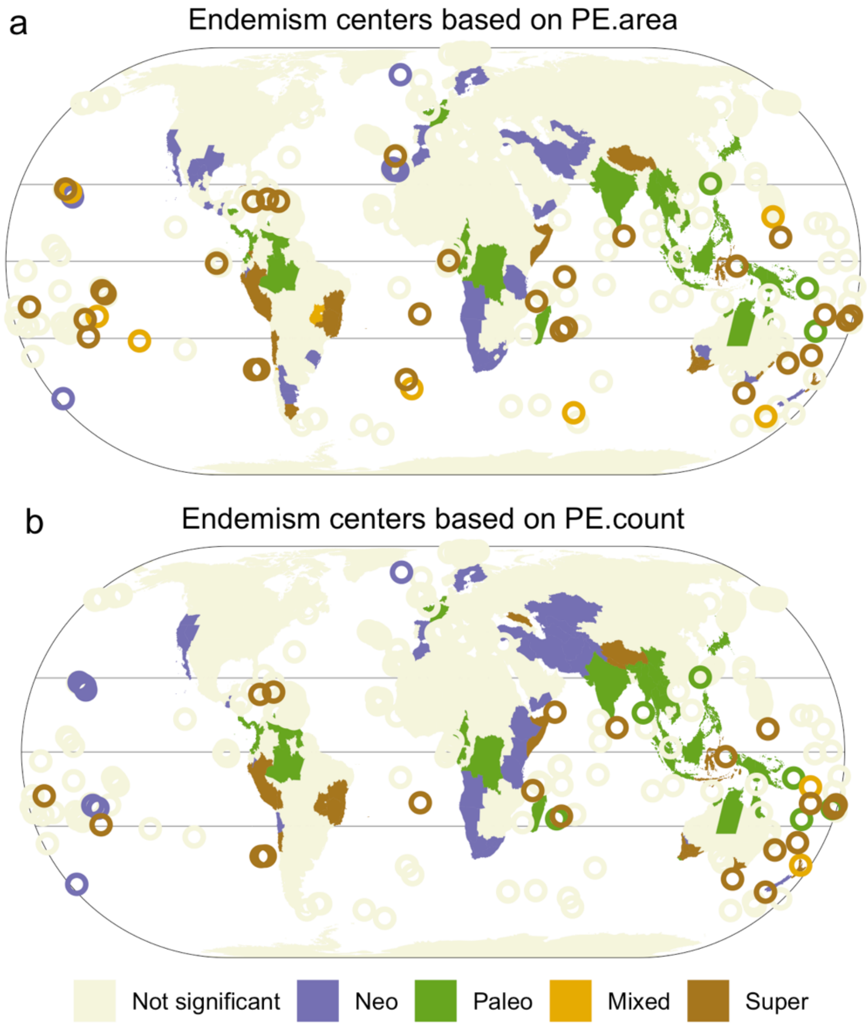


**Fig. S4** Global centers of neo- and paleoendemism for seed plants identified by the categorical analysis of neo- and paleoendemism (CANAPE) based on the distribution data including apomictic taxa. Colored regions present different types of endemism centers: violet, neoendemism; green, paleoendemism; yellow, mixed-endemism (i.e. neo- and paleoendemism); and brown indicating super-endemism (i.e. centers with both extremely high neo- and paleoendemism); beige, not significant. Patterns of neo- and paleoendemism are distinguished based on phylogenetic endemism with two competing ways of measuring species range size. PE.area indicates phylogenetic endemism calculated based on range size of species as the area of regions where a species occurs (a), while PE.count is calculated based on range size of species as the count of these regions (b).

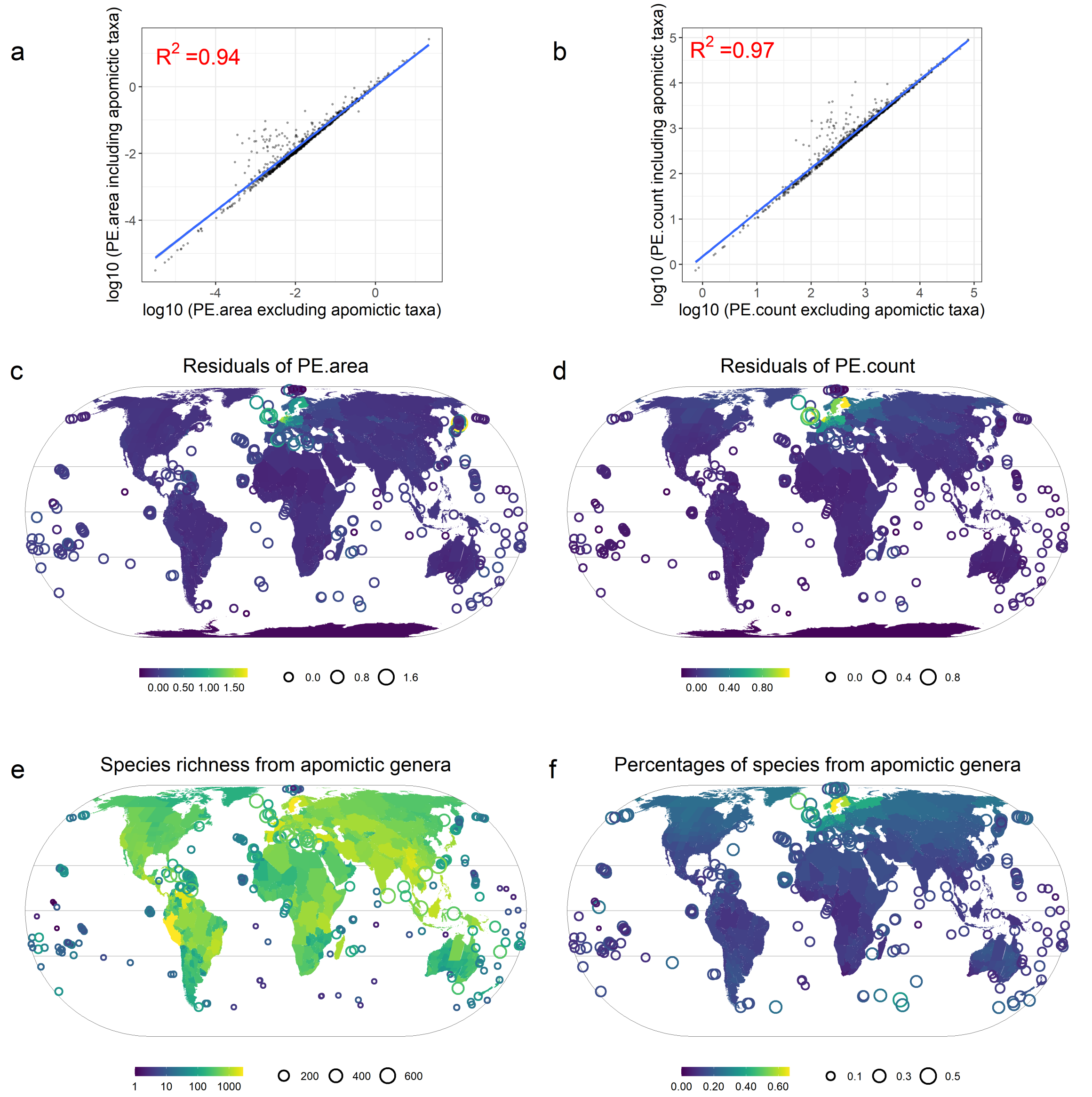


**Fig. S5** Comparison of phylogenetic endemism for seed plants based on the species distribution data including and excluding apomictic taxa. a and b, the linear regression between phylogenetic endemism based on the dataset including and excluding apomictic taxa; c and d, residuals from the linear regression. Phylogenetic endemism is calculated based on two different ways to measure range size of each species: in a and c, as the total area of regions a species occurs in (PE.area); and in b and d, as the number of these regions (PE.count). Positive residuals in c and d indicate higher values of phylogenetic endemism based on the data including apomictic taxa than expected based on the data excluding apomictic taxa. e, species richness of seed plants from genera known to include apomictic species. f, percentages of species from genera known to include apomictic species. $Log_{10}$ scale is used in e and all maps use Eckert IV projection.

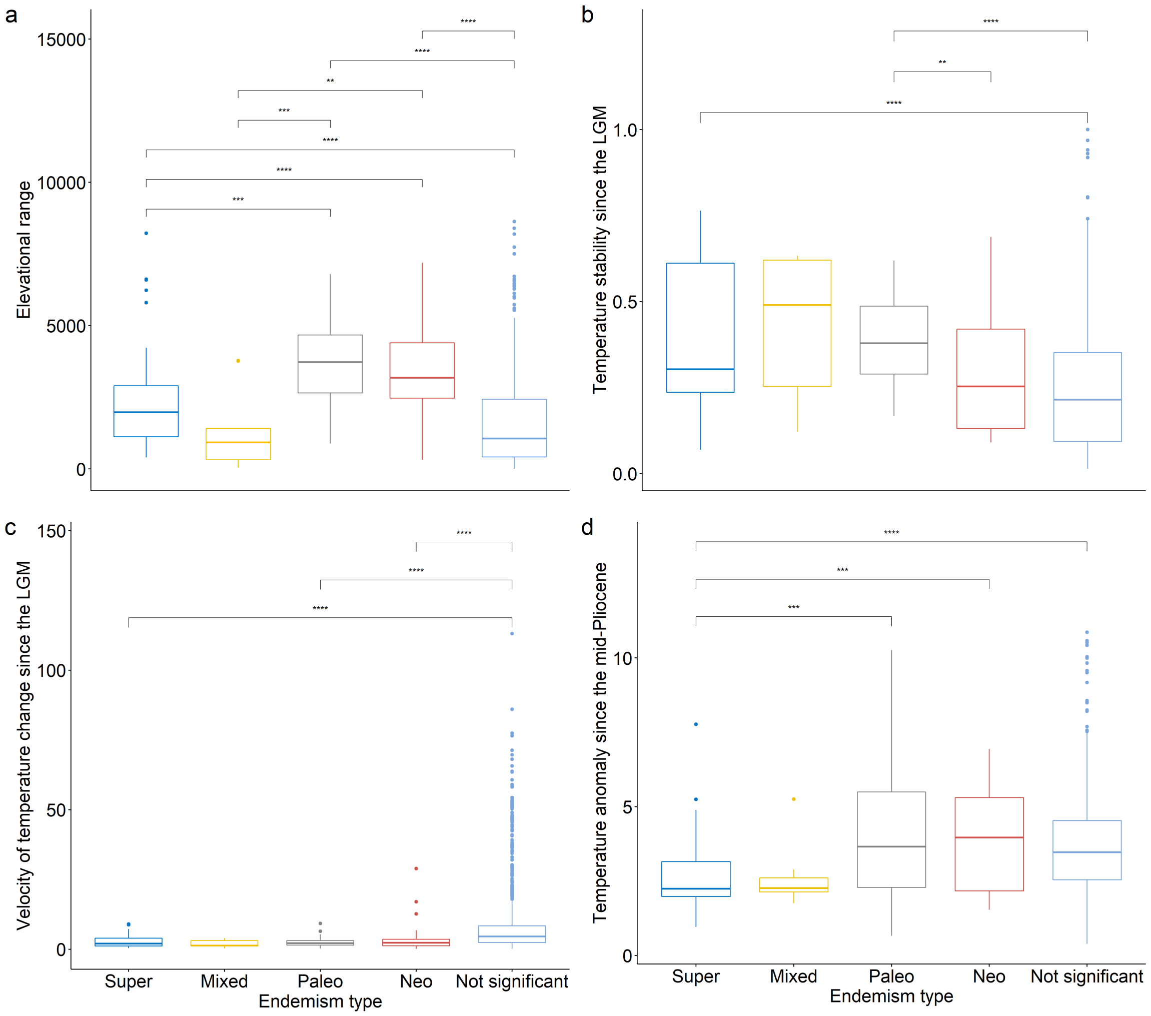


**Fig. S6** Distributions of environmental variables across regions of different endemism types based on phylogenetic endemism calculated using species range size as the area of regions where a species occurs. Endemism types are identified using the categorical analysis of neo- and paleoendemism (CANAPE) based on the distribution data excluding apomictic taxa. Distributions of environmental variables of the sampling regions are compared using pairwise Wilcoxon tests. Asterisks indicate statistical significance (*: p <= 0.05; **: p <= 0.01; ***: p <= 0.001; ****: p <= 0.0001). For each box, the middle horizontal line corresponds to the median; the lower and upper bounds of the box correspond to first and third quartiles, respectively. The upper whisker extends from the upper bound of the box to the highest value of the distribution, no further than 1.5 × interquartile range (i.e. distance between the first and third quartiles). The lower whisker extends from the lower bound of the box to the lowest value of the distribution, no further than 1.5 × interquartile range. Dots are values beyond the end of the whiskers ("outlier"). LGM = Last Glacial Maximum.

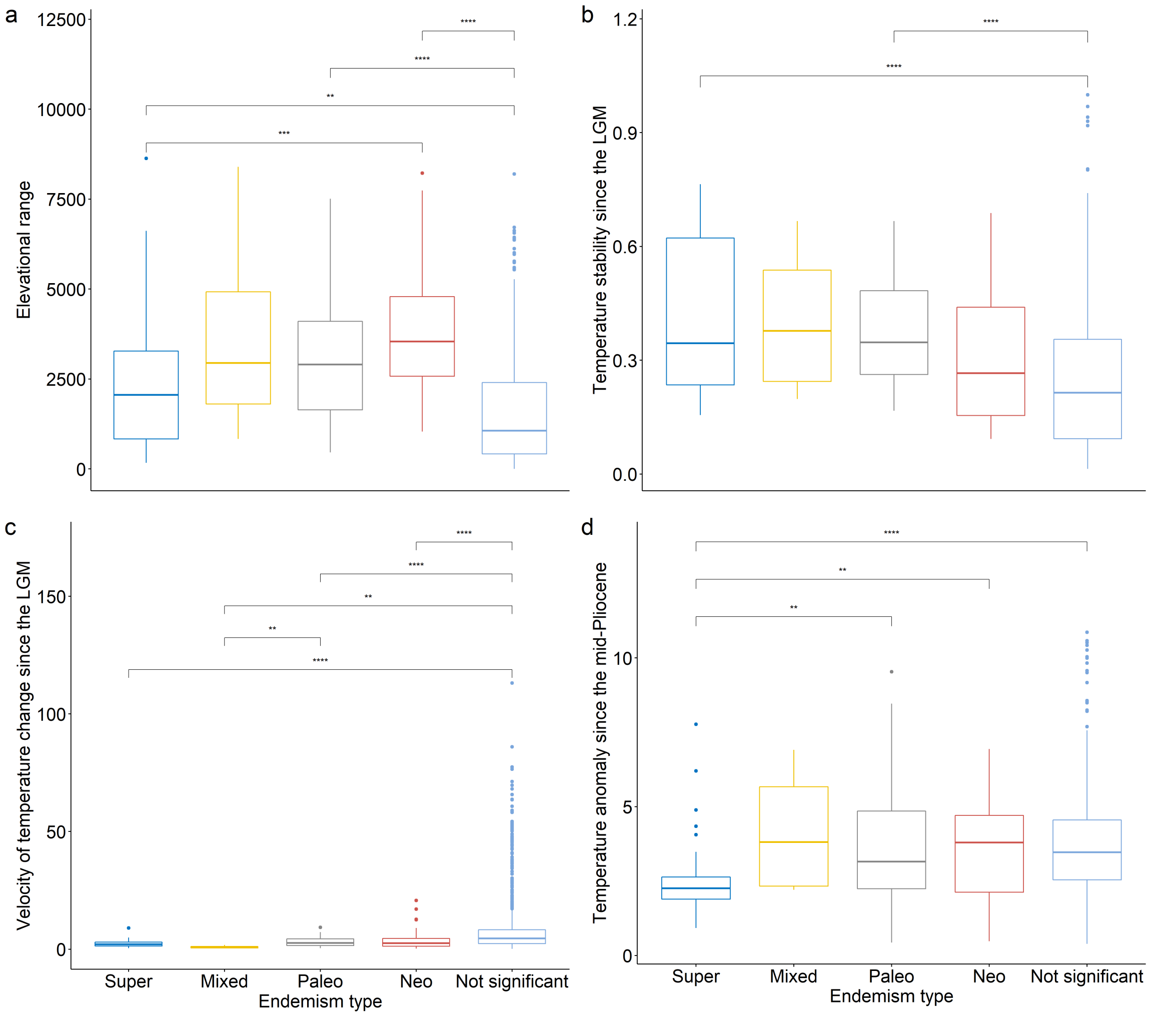


**Fig. S7** Distributions of environmental variables across regions of different endemism types based on phylogenetic endemism calculated using species range size as the number of regions where a species occurs. Endemism types are identified using the categorical analysis of neo- and paleoendemism (CANAPE) based on the distribution data excluding apomictic taxa. Distributions of environmental variables of the sampling regions are compared using pairwise Wilcoxon tests. Asterisks indicate statistical significance (*: p <= 0.05; **: p <= 0.01; ***: p <= 0.001; ****: p <= 0.0001). For each box, the middle horizontal line corresponds to the median; the lower and upper bounds of the box correspond to first and third quartiles, respectively. The upper whisker extends from the upper bound of the box to the highest value of the distribution, no further than 1.5 × interquartile range (i.e. distance between the first and third quartiles). The lower whisker extends from the lower bound of the box to the lowest value of the distribution, no further than 1.5 × interquartile range. Dots are values beyond the end of the whiskers ("outlier"). LGM = Last Glacial Maximum.

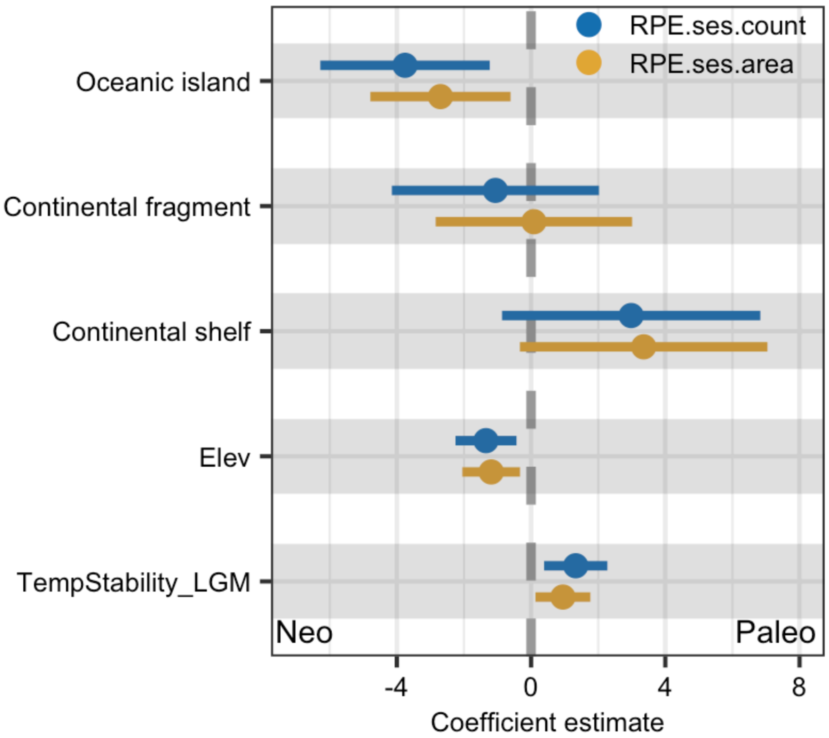


**Fig. S8** Determinants of neo- and paleoendemism based on the distribution data including apomictic taxa. Standardized regression coefficients of environmental factors are shown from spatial models of the standardized effect size of relative phylogenetic endemism of seed plants for regions with significantly high phylogenetic endemism. A positive effect of environmental factors in the model represents higher paleoendemism along the environmental factor, while a negative effect represents higher neoendemism. The reference level of geographic type is mainland regions. RPE.ses.area indicates the standardized effect size of relative phylogenetic endemism calculated based on range size of species as the area of regions where a species occurs, while RPE.ses.count is calculated based on range size of species as the count of these regions.

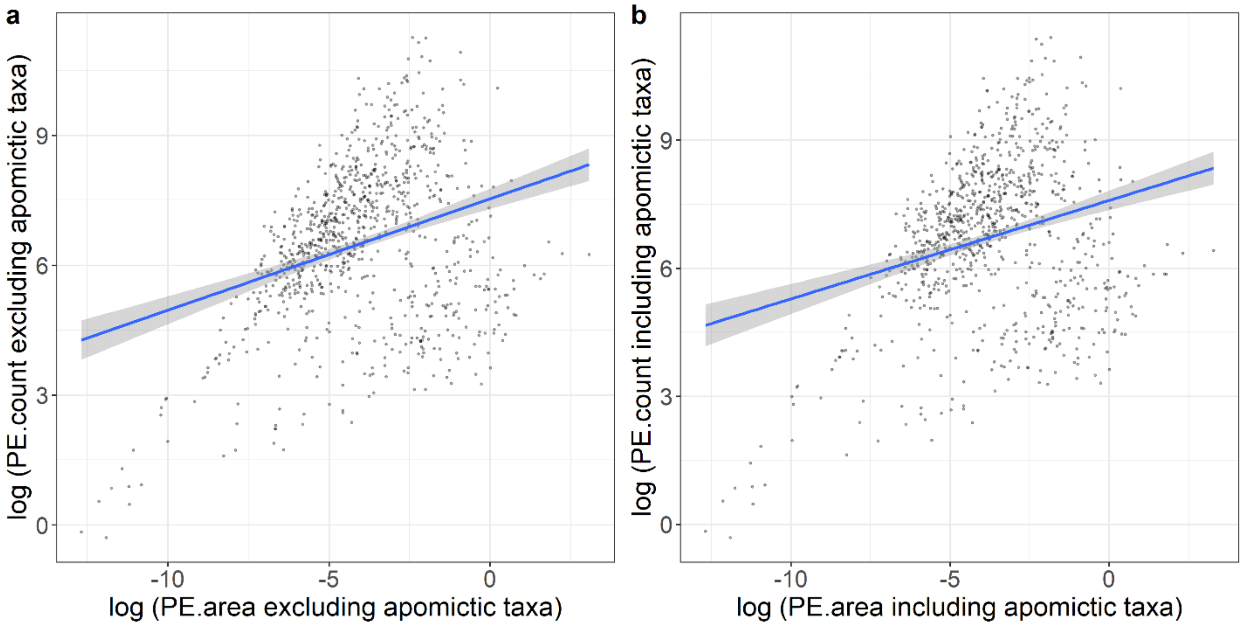


**Fig. S9** Comparison of phylogenetic endemism for seed plants with two competing ways of measuring species range size. PE.area indicates phylogenetic endemism calculated based on range size of species as the area of regions where a species occurs, while PE.count is calculated based on range size of species as the count of these regions. Phylogenetic endemism is calculated based on the dataset excluding (a) and including (b) apomictic taxa.

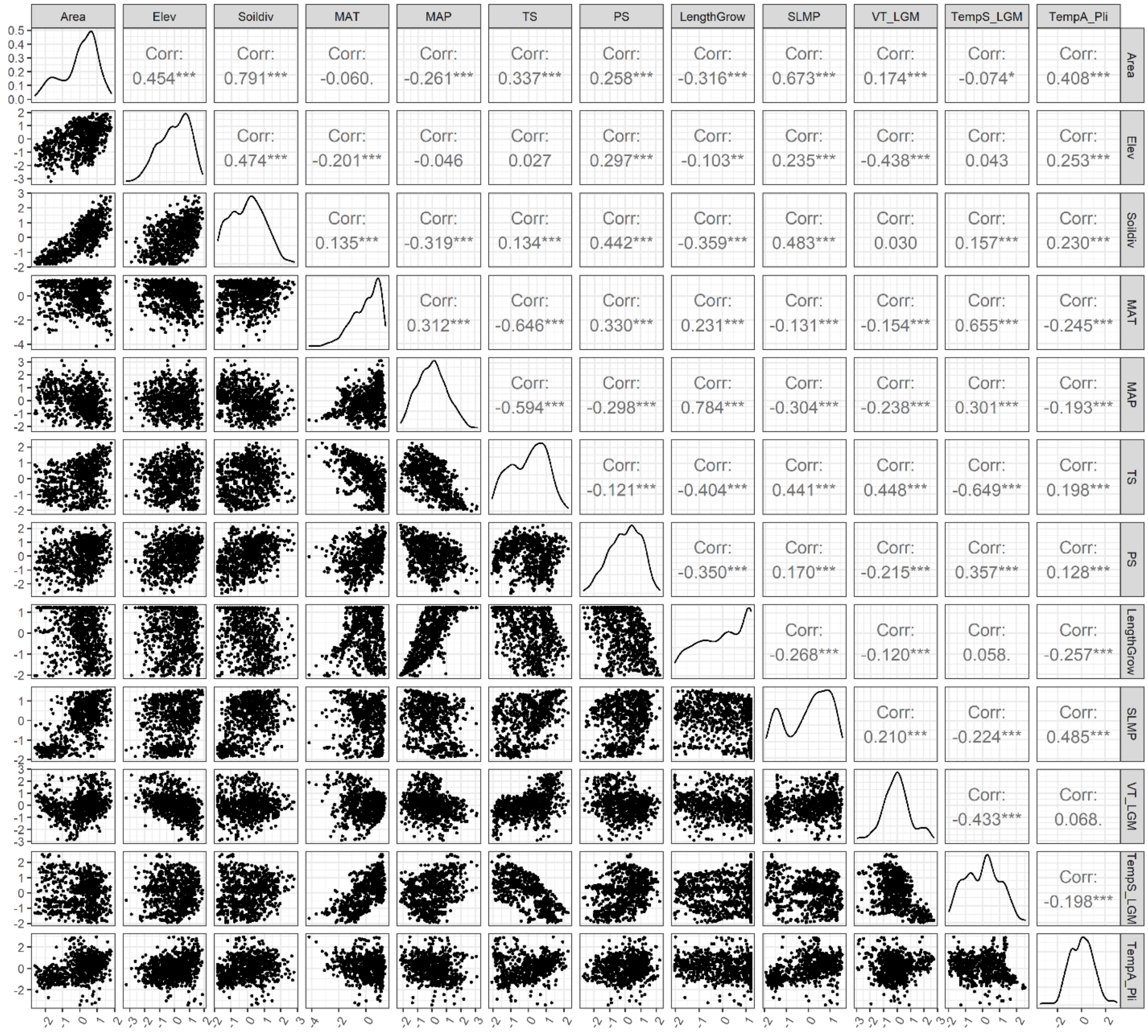


**Fig. S10** Correlations among all predictors and their density distributions after transformation. Numbers are Pearson correlation coefficients. Some predictors (i.e. Area, Elev, Soildiv, MAP, TS, PS, VT_LGM, TempS_LGM; TempA_Pli) are shown in log-scale as they were log-transformed for models owing to their skewed distributions. Area = region area; SLMP = surrounding landmass proportion; Elev = elevational range; Soildiv = number of soil types; MAT = mean annual temperature; MAP = mean annual precipitation; LengthGrow = length of growing season; TS = temperature seasonality; PS = precipitation seasonality; TempS_LGM = temperature stability since the Last Glacial Maximum; VT_LGM = velocity of temperature change since the Last Glacial Maximum; TempA_Pli = temperature anomaly between the mid-Pliocene warm period and present-day.

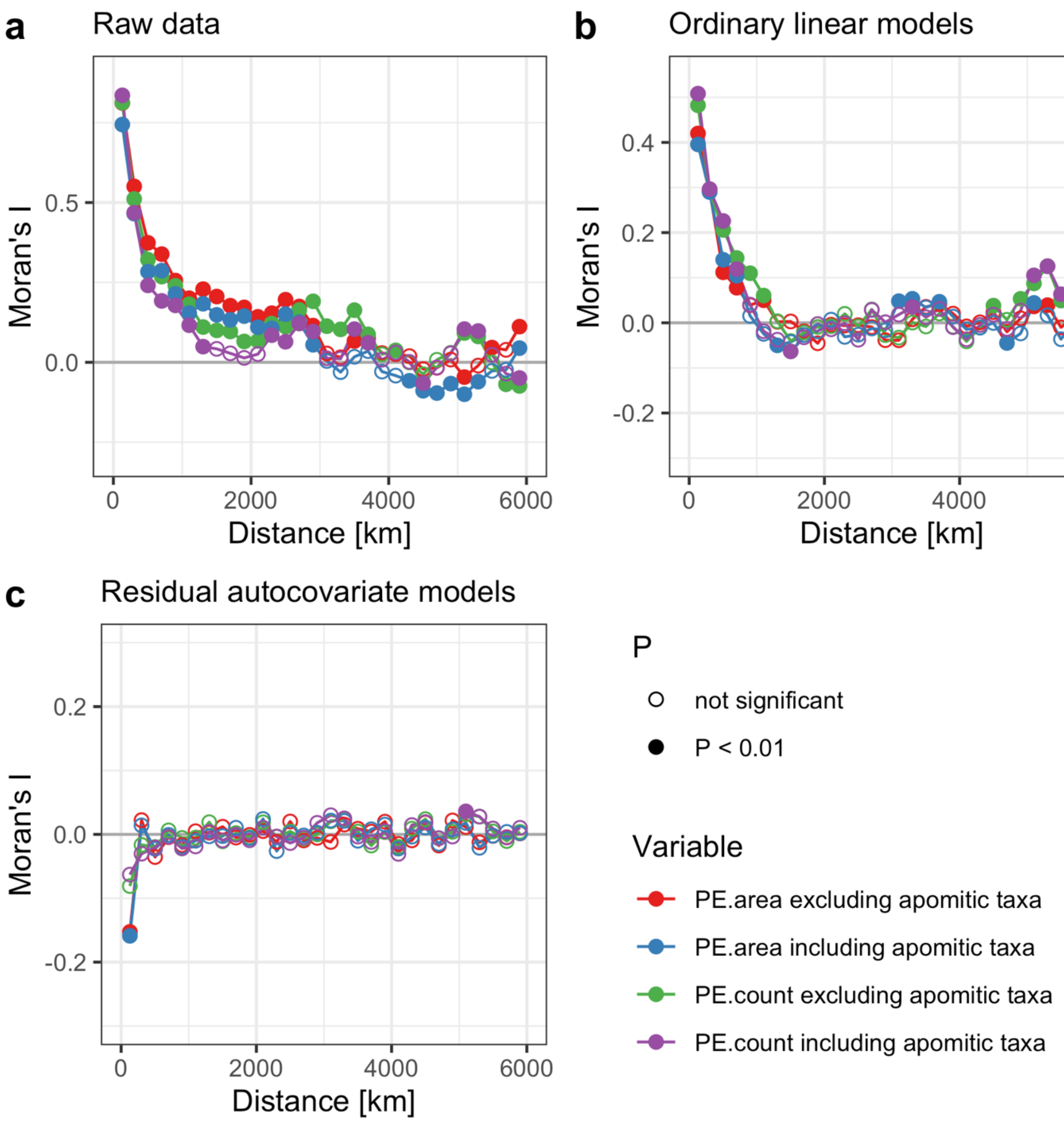


**Fig. S11** Spatial correlograms of raw phylogenetic endemism data (a), residuals from linear regression models for plant phylogenetic endemism (b) and from residual autocovariate models (c). Full symbols indicate a significant Moran's I at a given lag distance (P < 0.01). PE.area indicates phylogenetic endemism calculated based on species range size as the area of regions where a species occurs, while PE.count is calculated based on species range size as the count of these regions.

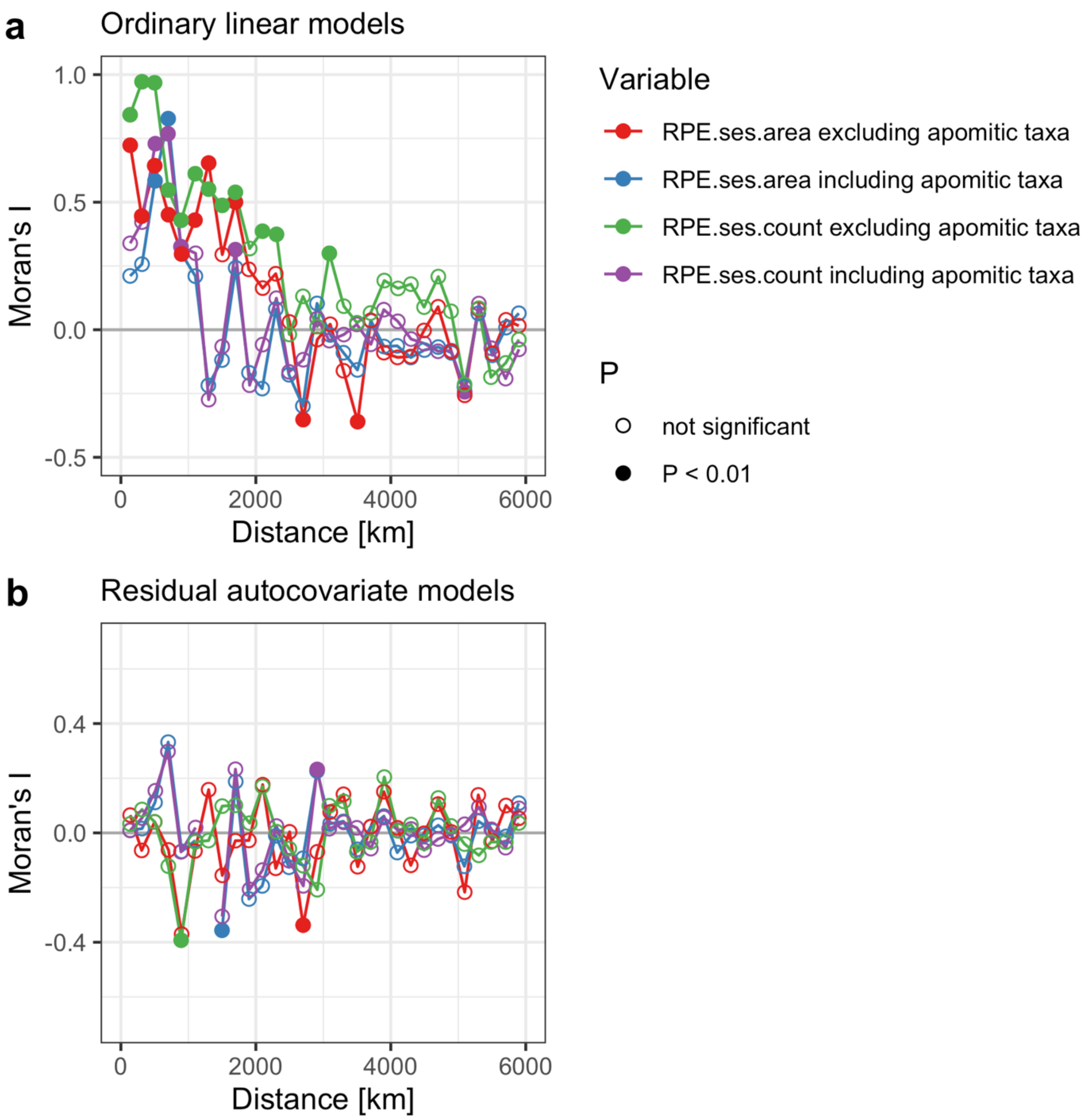


**Fig. S12** Spatial correlograms of residuals for standardized effect size of relative phylogenetic endemism from linear regression models (a) and from residual autocovariate models (b). Full symbols indicate a significant Moran's I at a given lag distance (P < 0.01). RPE.ses.area indicates standardized effect size of relative phylogenetic endemism calculated based on range size of species as the area of regions where a species occurs, while RPE.ses.count is calculated based on range size of species as the count of these regions.

**Table S1** Hypotheses and related predictors of phylogenetic endemism in plants. ↑ represents effects on phylogenetic endemism hypothesized to be positive, while ↓ represents negative effects.

| Factor | Variables | Hypothesis |
|---|---|---|
| Isolation | Surrounding landmass proportion (↓); whether or not a region is an island | Isolation fosters allopatric speciation by reducing gene flow, which in turn promotes endemism (3, 4). |
| Environmental heterogeneity | Elevational range (↑); number of soil types (↑) | Heterogeneous regions include small habitats supporting more narrow-ranged species, allowing for geographic isolation promoting specialization, and serving as refugia during unfavorable climate change periods (5, 6). |
| Energy and water availability | Mean annual temperature (↑); mean annual precipitation (↑); length of growing season (↑) | Warm and humid climates are hypothesized to support larger population sizes in small regions by offering sufficient resources, which promotes long-term survival of spatially restricted lineages and their accumulation over long timescales (7). Additionally, high energy availability may increase the opportunity for speciation, which promotes endemism (8, 9). |
| Climatic seasonality | Temperature seasonality (↓); precipitation seasonality (↓) | High climatic seasonality selects for species with broader climatic tolerances and large ranges (10). |
| Long-term climatic stability | Temperature stability since the Last Glacial Maximum (LGM) (↑); velocity of temperature change since the LGM (↓); temperature anomaly since the mid-Pliocene warm period (↓) | Long-term climatic stability allows for the evolution of narrow physiological tolerances and specialization, and reduces extinction risk of small-ranged lineages (11–13) |

**Table S2** Hypotheses and related predictors of neo- and paleoendemism in plants.

| **Factor** | **Variables** | **Hypotheses** |
|---|---|---|
| Geologic origins | Geographic type of a region based on their geologic origins, i.e. continental shelf islands, continental fragments, oceanic islands, or mainland. | Oceanic islands host higher neoendemism, while continental fragments are centers of paleoendemism (14). |
| Montane regions | Elevational range | On the one hand, montane regions are centers of neoendemism, due to high speciation rates that are driven by long-term orogenic and climatic dynamics in mountains (15, 16); on the other hand, montane regions foster paleoendemism, because montane regions promote the persistence of ancient lineages during unfavorable climate change periods (17). The two processes may be not mutually exclusive, leading to montane regions as centers of both neo- and paleoendemism (18). |
| Past climate change | Temperature stability since the Last Glacial Maximum (LGM); velocity of temperature change since the LGM; temperature anomaly since the mid-Pliocene warm period | Regions with stable climates have suffered less severe environmental changes across space and may have acted as refugia where plants could persist over time, and host higher paleoendemism (19). |

**Table S3** Top ten islands of different minimum size with highest phylogenetic endemism for seed plants based on the distribution data excluding apomictic taxa. PE.area indicates phylogenetic endemism calculated based on range size of each species measured as the total area of regions a species occurs in.

| island >100 km$^2$ | PE.area | island >1000 km$^2$ | PE.area |
|---|---|---|---|
| Mahe | 2.92 | Caroline Islands | 1.93 |
| St. Helena | 2.06 | New Caledonia | 1.27 |
| Caroline Islands | 1.93 | Mauritius | 1.10 |
| Raiatea | 1.37 | Kaua'i Island | 0.96 |
| Rodrigues | 1.27 | Tahiti | 0.83 |
| New Caledonia | 1.27 | La Réunion | 0.77 |
| Mauritius | 1.10 | O'ahu Island (incl. Mokoli'i Islet) | 0.66 |
| Principe | 1.08 | Samoa | 0.60 |
| Sao Tomé | 1.02 | Tenerife | 0.56 |
| Madeira | 0.96 | Jamaica | 0.56 |

**Table S4** Top ten regions of phylogenetic endemism for seed plants based on the distribution data including apomictic taxa. Phylogenetic endemism is calculated based on two different ways to measure range size of each species: (i) as the total area of regions a species occurs in (PE.area) and (ii) as the number of these regions (PE.count).

| PE.area | | | | | | | | PE.count | | | |
|---|---|---|---|---|---|---|---|---|---|---|---|
| **Mainland** | **PE** | **island >10 $km^2$** | **PE** | **island >100 $km^2$** | **PE** | **island >1000 $km^2$** | **PE** | **Mainland** | **PE** | **Island** | **PE** |
| Costa Rica | 0.49 | Lord Howe Island | 26.56 | Mahe | 3.03 | Caroline Islands | 2.10 | Peru | 86902 | Madagascar | 90040 |
| Pichincha, Ecuador | 0.47 | Masatierra | 10.13 | St. Helena | 2.33 | Mauritius | 1.44 | Western Cape Province, South Africa | 56603 | Borneo | 76810 |
| Panama | 0.43 | Tubuai Isand | 6.21 | Caroline Islands | 2.10 | New Caledonia | 1.43 | Venezuela | 51928 | Papua New Guinea | 55351 |
| Western Cape Province, South Africa | 0.41 | Norfolk Island | 6.01 | Mauritius | 1.44 | La Réunion | 1.40 | Vietnam | 37975 | Philippines | 51794 |
| Carchi, Ecuador | 0.32 | Masafuera | 5.64 | New Caledonia | 1.43 | Kaua'i Island | 1.05 | Thailand | 34753 | Indonesian New Guinea | 31787 |
| Valle del Cauca, Colombia | 0.26 | Silhouette | 4.04 | Raiatea | 1.40 | Tahiti | 0.84 | India | 34545 | Sumatra | 29958 |
| Rio de Janeiro, Brazil | 0.26 | Mahe | 3.03 | La Réunion | 1.40 | Samoa | 0.75 | Asiatic Turkey | 34509 | Cuba | 27660 |
| Antioquia, Colombia | 0.25 | Rarotonga | 2.54 | Rodrigues | 1.36 | O'ahu Island (incl. Mokoli'i Islet) | 0.75 | Yunnan | 33386 | New Caledonia | 27282 |
| Quindío, Colombia | 0.25 | St. Helena | 2.33 | Madeira | 1.13 | Comoros | 0.73 | Minas Gerais | 32888 | Japan | 21096 |
| Loja, Ecuador | 0.24 | Henderson | 2.30 | Principe | 1.08 | Tenerife | 0.67 | Panama | 31249 | Hispaniola | 17330 |

**Table S5** Linear model results for phylogenetic endemism of seed plants. Models are fitted for phylogenetic endemism based on two different methods used to quantify species range size and distribution data excluding and including apomictic taxa. The model performances are quantified by Akaike's information criterion (AIC) and adjusted coefficient of determination ($R^2$). PE.area indicates phylogenetic endemism calculated based on range size of species as the area of regions where a species occurs, while PE.count is calculated based on range size of species as the count of these regions. Area = region area; SLMP = surrounding landmass proportion; Elev = elevational range; Soildiv = number of soil types; MAT = mean annual temperature; MAP = mean annual precipitation; LengthGrow = length of growing season; TS = temperature seasonality; PS = precipitation seasonality; TempStability_LGM = temperature stability since the Last Glacial Maximum; VT_LGM = velocity of temperature change since the Last Glacial Maximum; TempAnomaly_midPliocene = temperature anomaly between the mid-Pliocene warm period and present-day; Geo_class = whether or not a region is an island. The rest terms are the interactions between each predictors and SLMP (or Geo_class).

| Models | Dataset for modeling excluding apomictic taxa | | | | Dataset for modeling including apomictic taxa | | | |
|---|---|---|---|---|---|---|---|---|
| | PE.area | | PE.count | | PE.area | | PE.count | |
| | AIC | $R^2$ | AIC | $R^2$ | AIC | $R^2$ | AIC | $R^2$ |
| Area + SLMP + Elev + Soildiv + MAT + MAP + LengthGrow + TS + PS + VT_LGM + TempStability_LGM + TempAnomaly_midPliocene + Area:SLMP + Elev:SLMP + Soildiv:SLMP + MAT:SLMP + MAP:SLMP + LengthGrow:SLMP + TS:SLMP + PS:SLMP + VT_LGM:SLMP + TempStability_LGM:SLMP + TempAnomaly_midPliocene:SLMP | 1197.439 | 0.658 | 524.456 | 0.786 | 1169.485 | 0.635 | 460.578 | 0.791 |
| Area + Geo_class + Elev + Soildiv + MAT + MAP + LengthGrow + TS + PS + VT_LGM + TempStability_LGM + TempAnomaly_midPliocene + Area:Geo_class + Elev:Geo_class + Soildiv:Geo_class + MAT:Geo_class + MAP:Geo_class + LengthGrow:Geo_class +TS:Geo_class + PS:Geo_class + VT_LGM:Geo_class + TempStability_LGM:Geo_class + TempAnomaly_midPliocene:Geo_class | 1226.713 | 0.645 | 551.191 | 0.779 | 1194.351 | 0.624 | 490.216 | 0.784 |

**References S1** References of checklists and floras from the Global Inventory of Floras and Traits (GIFT) used to compile the regional species composition data.


1. K. F. Kenneally, Ashmore Reef and Cartier Island: Species lists.
2. Christmas Island National Park, Third Christmas Island national park management plan (Parks Australia North, 2002).
3. D.J. Du Puy, Christmas Island: species lists.
4. P.S. Green, Lord Howe Island: species lists.
5. P.S. Green, Norfolk Island: Species lists.
6. R.J. Hnatiuk, Subantarctic Islands: species lists.
7. M. Arechavaleta, S. Rodríguez, N. Zurita, A. García, Lista de especies silvestres de Canarias. Hongos, plantas y animales terrestres (Consejería de Medio Ambiente y Ordenación Territorial, Gobierno de Canarias, 2009).
8. D. A. Broughton, J. H. McAdam, A checklist of the native vascular flora of the Falkland Islands (Islas Malvinas): new information on the species present, their ecology, status and distribution. The Journal of the Torrey Botanical Society 132, 115–148 (2005).
9. W.L. Wagner, D.R. Herbst, D.H. Lorence, Flora of the Hawaiian Islands website.
10. H. Kristinsson, Checklist of the vascular plants of Iceland (Náttúrufræðistofnun Íslands, 2008).
11. A. C. Robinson, P. D. Canty, D. Fotheringham, Investigator group expedition 2006: flora and vegetation. Trans. R. Soc. S. Aust. 132, 173–220 (2008).
12. F. Kirchner, F. Picot, E. Merceron, G. Gigot, Flore vasculaire de La Réunion (Conservatoire Botanique National de Mascarin, 2010).
13. W. L. Wagner, D. H. Lorence, Flora of the Marquesas Islands website.
14. N. Kingston, S. Waldren, U. Bradley, The phytogeographical affinities of the Pitcairn Islands – a model for south-eastern Polynesia? Journal of Biogeography 30, 1311–1328 (2003).
15. J. S. Releford, J. Stevens, K. W. Bridges, W. C. McClatchey, Flora of Rongelap and Ailinginae Atolls, Republic of the Marshall Islands. Atoll Research Bulletin 572, 1–13 (2009).
16. K. Nakamura, R. Suwa, T. Denda, M. Yokota, Geohistorical and current environmental influences on floristic differentiation in the Ryukyu Archipelago, Japan. Journal of Biogeography 36, 919–928 (2009).
17. B. E. Sandbakk, I. G. Alsos, G. Arnesen, R. Elven, The flora of Svalbard.
18. S. A. Renvoize, A floristic analysis of the western Indian Ocean coral islands. Kew Bulletin 30, 133–152 (1975).
19. P. Acevedo-Rodríguez, M. T. Strong, Catalogue of the seed plants of the West Indies Website.

20. R. Taylor, Straight through from London: the Antipodes and Bounty Islands, New Zealand (Heritage Expeditions New Zealand, 2006).
21. N. M. Wace, The vegetation of Gough Island. Ecological Monographs 31, 337–367 (1961).
22. A.G. Miller, M. Morris, Ethnoflora of the Soqotra Archipelago (Royal Botanic Garden, 2004).
23. F.R. Fosberg, S.A. Renvoize, C.C. Townsend, The flora of Aldabra and neighbouring islands (HMSO, 1980).
24. W.R. Sykes, Contributions to the flora of Niue. Bulletin. Department of Scientific and Industrial Research, New Zealand 200, 321 (1970).
25. W.R. Sykes, C. J. West, J. E. Beever, A. J. Fife, Kermadec Islands flora-special edition: a compilation of modern material about the flora of the Kermadec Islands (Manaaki Whenua Press, Landcare Research, 2000).
26. A.W. Exell, Catalogue of the vascular plants of S. Tome (with Principe and Annobon) (Trustees of the British Museum, 1944).
27. P. Ashmole, M. Ashmole, St Helena and Ascension Island: A natural history (Anthony Nelson Ltd, 2000).
28. T. G. Yuncker, Plants of Tonga. Bishop Museum Bulletin 220, 1–283 (1959).
29. CARMABI, Dutch Caribbean Biodiversity Explorer.
30. N. M. Wace, J. H. Dickson, The terrestrial botany of the Tristan da Cunha Islands. Philosophical Transactions of the Royal Society of London B Biological Sciences 249, 273–360 (1965).
31. S. W. Greene, D. W.H. Walton, An annotated check list of the sub-Antarctic and Antarctic vascular flora. Polar Record 17, 473–484 (1975).
32. Q. C. B. Cronk, The past and present vegetation of St Helena. J Biogeogr 16, 47–64 (1989).
33. N. Vander Velde, The vascular plants of Majuro atoll, Republic of the Marshall Islands. Atoll Research Bulletin 503, 1–141 (2003).
34. H. Takahashi, V. Y. Barkalov, S. Gage, Y.N. Zhuravlev, A preliminary study of the flora of Chirpoi, Kuril Islands. J. Hattori Bot. Lab. 48, 31–42 (1997).
35. M. Charters, Flora of Bermuda.
36. P.A.V. Borges, C. Abreu, A.M.F. Aguiar, P. Carvalho, R. Jardim, I. Melo, P. Oliveira, C. Sérgio, A.R.M. Serrano, P. Vieira, Listagem dos fungos, flora e fauna terrestres dos arquipélagos da Madeira e Selvagens (Direcção Regional do Ambiente da Madeira and Universidade dos Açores, 2008).
37. M. Arechavaleta, N. Zurita, M.C. Marrero, J.L. Martín, Lista preliminar de especies silvestres de Cabo Verde (hongos, plantas y animales terrestres) (Consejería de Medio Ambiente y Ordenación Territorial, Gobierno de Canarias, 2005).
38. J. Florence, H. Chevillotte, C. Ollier, J.-Y. Meyer, Base de données botaniques Nadeaud de l'Herbier de la Polynésie française (PAP).

39. K. Y. Chong, T.W.H. Tan, R. T. Corlett, A checklist of the total vascular plant flora of Singapore: Native, Naturalised and Cultivated Species (Raffles Museum of Biodiversity Research, 2009).
40. UIB, Herbario virtual del Mediterráneo Occidental.
41. P. N. Johnson, D. J. Campbell, Vascular plants of the Auckland Islands. New Zealand journal of botany 13, 665–720 (1975).
42. B.R. Jackes, Plants of Magnetic Island (James Cook University, 2010).
43. C. Marticorena, T. F. Stuessy, C. M. Baeza, Catalogue of the vascular flora of the Robinson Crusoe or Juan Fernández islands, Chile. Gayana Botánica 55, 187–211 (1998).
44. M.-H. Sachet, Flora and vegetation of Clipperton Island. Proceedings of the California Academy of Sciences 31, 249–307 (1962).
45. S. Gage, S.L. Joneson, V. Y. Barkalov, N.A. Eremenko, H. Takahashi, A newly compiled checklist of the vascular plants of the Habomais, the Little Kurils. Bulletin of the Hokkaido University Museum 3, 67–91 (2006).
46. M.L. Baker, M.F. Duretto, A census of the vascular plants of Tasmania (Tasmanian Herbarium, Tasmanian Museum and Art Gallery, 2011).
47. H. Takahashi, V. Y. Barkalov, S. Gage, B. Semsrott, M. Ilushko, Y.N. Zhuravlev, A floristic study of the vascular plants of Kharimkotan, Kuril Islands. Bulletin of the Hokkaido University Museum 3, 41–66 (2006).
48. M. Tatewaki, Geobotanical studies on the Kurile Islands. Acta Horti Gotoburgensis 21, 43–123 (1957).
49. R.R. Thaman, F.R. Fosberg, H.I. Manner, D.C. Hassall, The flora of Nauru. Atoll Research Bulletin 392, 1–233 (1994).
50. W. McClatchey, R. Thaman, S. Vodonaivalu, A preliminary checklist of the flora of Rotuma with Rotuman names. Pacific Science 54, 345–363 (2000).
51. P. J. de Lange, P. B. Heenan, J. R. Rolfe, Checklist of vascular plants recorded from Chatham Islands (2011).
52. J. Searle, S. Madden, Flora assessment of South Stradbroke Island (Gold Coast City Council, 2006).
53. T. Abe, Threatened pollination systems in native flora of the Ogasawara (Bonin) Islands. Annals of Botany (London) 98, 317 (2006).
54. C. M. Cheffings, L. Farrell, Eds., The vascular plant red data list for Great Britain (Joint Nature Conservation Committee, 2005).
55. G. V. Byrd, Vascular vegetation of Buldir Island, Aleutian Islands, Alaska, compared to another Aleutian Island. Arctic 37, 37–48 (1984).
56. W. G. D'Arcy, The island of Anegada and its flora. Atoll Research Bulletin 139, 1–21 (1971).
57. F.R. Fosberg, M.H. Sachet, Flora of Maupiti, Society Islands. Atoll Research Bulletin 294, 1–70 (1987).

58. D.R. Stoddart, F.R. Fosberg, Flora of the Phoenix Islands, central Pacific. Atoll Research Bulletin 393, 1–60 (1994).
59. L. Wester, Checklist of the vascular plants of the northern Line Islands. Atoll Research Bulletin 187, 1–38 (1985).
60. H. St John, Flora of Eniwetok Atoll. Pacific Science 14, 313–336 (1960).
61. E.D. Marquand, Flora of Guernsey and the lesser Channel Islands: namely Alderney, Sark, Herm, Jethou, and the adjacent islets (Dulau & Co, 1901).
62. L.V. Lester-Garland, A flora of the islands of Jersey: with a list of the plants of the Channel Islands in general, and remarks upon their distribution and geographical affinities (West, Newman & Co, 1903).
63. L. Sáez, J. A. Rosselló, Llibre vermell de la flora vascular de les Illes Balears (Direcció General de Biodiversitat, Conselleria de Medi Ambient, Govern de les Illes Balears, 2001).
64. C.A. Stace, R.G. Ellis, D.H. Kent, D.J. McCosh, Vice-county Census Catalogue of the vascular plants of Great Britain, the Isle of Man and the Channel Islands (Botanical Society of the British Isles, 2003).
65. M. Kerguelen, Base de Données Nomenclaturales de la Flore de France.
66. R. Jahn, P. Schönfelder, Exkursionsflora für Kreta (Ulmer (Eugen), 1995).
67. F. Conti, G. Abbate, A. Alessandrini, C. Blasi, Annotated Checklist of the Italian Vascular Flora (Palombi Editori, 2005).
68. University of Kent, Cook Islands Biodiversity and Ethnobiology Database.
69. J. D. Shaw, D. Spear, M. Greve, S. L. Chown, Taxonomic homogenization and differentiation across Southern Ocean Islands differ among insects and vascular plants. J Biogeogr 37, 217–228 (2010).
70. A.M. Chernyaeva, Flora of Onekotan Island. Bulletin of Main Botanical Garden 87, 21–29 (1973).
71. E.M. Egorova, Flora of Shiashkotan Island. Bulletin of the Main Botanical Garden 54, 114–120 (1964).
72. P. Morat, T. Jaffré, F. Tronchet, J. Munziger, Y. Pillon, J.-M. Veillon, M. Chalopin, P. Birnbaum, F. Rigault, G. Dagostini, J. Tinerl, P.P. Lowry II, The taxonomic reference base "Florical" and characteristics of the native vascular flora of New Caledonia. Adansonia 34, 179–221 (2012).
73. J. Gerlach, The biodiversity of the granitic islands of Seychelles. Phelsuma 11 (Supplement A), 1–47 (2003).
74. L. Raulerson, Checklist of plants of the Mariana Islands. University of Guam Herbarium Contribution 37, 1–69 (2006).
75. P.A.V. Borges, A. Costa, R. Cunha, R. Gabriel, V. Gonçalves, A. F. Martins, I. Melo, M. Parente, P. Raposeiro, P. Rodrigues, R. S. Santos, L. Silva, P. Vieira, V. Vieira, A list of the terrestrial and marine biota from the Azores (Princípia, 2010).

76. C. T. Imada, Ed., Hawaiian Native and Naturalized Vascular Plants Checklist (Hawaii Biological Survey, Bishop Museum, 2012).
77. A. C. Smith, Flora Vitiensis nova: a new Flora of Fiji (spermatophytes only) (Pacific Tropical Botanical Garden, 1979-1996).
78. W. A. Strahm, The conservation and restoration of the flora of Mauritius and Rodrigues.
79. J. P. Price, W. L. Wagner, A phylogenetic basis for species–area relationships among three Pacific Island floras. American Journal of Botany 98, 449–459 (2011).
80. C. Costion, D. Lorence, The endemic plants of Micronesia: A geographical checklist and commentary. Micronesica 43, 51–100 (2012).
81. 3D Environmental, Vegetation Communities and Regional Ecosystems of The Torres Strait Islands, Queensland, Australia (2008).
82. R. P. Pandey, P. G. Diwakar, An integrated checklist flora of Andaman and Nicobar Islands, India. J. Econ. Taxon. Bot 32, 403–500 (2008).
83. A. Gray, T. Pelembe, S. Stroud, The conservation of the endemic vascular flora of Ascension Island and threats from alien species. Oryx 39, 449–453 (2005).
84. E. Figueiredo, J. Paiva, T. Stevart, F. Oliveira, G. F. Smith, Annotated catalogue of the flowering plants of São Tomé and Príncipe. Bothalia 41, 41–82 (2011).
85. M. Velayos, P. Barberá, F. J. Cabezas, M. d. La Estrella, M. Fero, C. Aedo, Checklist of the Vascular Plants of Annobón (Equatorial Guinea). Phytotaxa 171, 1–78 (2014).
86. Tela Botanica, Base de données des Trachéophytes de France métropolitaine (bdtfx).
87. E. K. Cameron, P. J. de Lange, J. McCallum, G. A. Taylor, P. J. Bellingham, Vascular flora and some fauna for a chain of six Vascular flora and some fauna for a chain of six Hauraki Gulf islands east and southeast of Waiheke Island. Auckland Botanical Society Journal 62, 136–156 (2007).
88. E. A. Brown, Vegetation and flora of Ponui Island, Hauraki Gulf, New Zealand. TANE 25, 5–16 (1979).
89. W. R. Barker, R. M. Barker, J. P. Jessop, H. P. Vonow, Census of South Australian vascular plants. Journal of the Adelaide Botanic Gardens Supplement 1, 1–396 (2005).
90. BioScripts, Flora Vascular.
91. G. Brundu, I. Camarda, The Flora of Chad: a checklist and brief analysis. Phytokeys 23, 1–17 (2013).
92. J. de Egea, M. Peña-Chocarro, C. Espada, S. Knapp, Checklist of vascular plants of the Department of Ñeembucú, Paraguay. Phytokeys 9, 15–179 (2012).
93. V. A. Funk, T. Hollowell, P. Berry, C. Kelloff, S. N. Alexander, Checklist of the plants of the Guiana Shield (Venezuela: Amazonas, Bolivar, Delta Amacuro; Guyana, Surinam, French Guiana) (Department of Botany, National Museum of Natural History, 2007).

94. ZDSF, SKEW, Info Flora: Artenliste Schweiz 5x5 km.
95. F. O. Zuloaga, O. Morrone, M. Belgrano, Catálogo de las Plantas Vasculares del Cono Sur.
96. C. Marticorena, F. A. Squeo, G. Arancio, M. Muñoz, "Catálogo de la flora vascular de la IV Región de Coquimbo" in Libro rojo de la flora nativa y de los sitios prioritarios para su conservación: Región de Atacama, F. A. Squeo, G. Arancio, J. R. Gutiérrez, Eds. (Ediciones Universidad de La Serena La Serena, 2008), pp. 105–142.
97. J. Norton, S. A. Majid, D. Allan, M. Al Safran, B. Böer, R. A. Richer, An illustrated checklist of the flora of Qatar (Browndown Publications Gosport, 2009).
98. Queensland Government, Census of the Queensland flora 2014.
99. SLUFG, Rote Liste und Artenliste Sachsen: Farn- und Samenpflanzen (Sächsisches Landesamt für Umwelt, Landwirtschaft und Geologie, 2015).
100. SANBI, Plants of Southern Africa: An online checklist.
101. P. S. Short, D. E. Albrecht, I. D. Cowie, D. L. Lewis, B. M. Stuckey, Checklist of the vascular plants of the Northern Territory (Department of Natural Resources, Environment, The Arts and Sport, 2011).
102. USDA, NRCS, The PLANTS Database.
103. C. Vogt, Composición de la Flora Vascular del Chaco Boreal, Paraguay. I. Pteridophyta y Monocotiledoneae. Steviana 3, 13–47 (2011).
104. L. Catarino, E. S. Martins, M. F. Basto, M. A. Diniz, An annotated checklist of the vascular flora of Guinea-Bissau (West Africa). Blumea-Biodiversity, Evolution and Biogeography of Plants 53, 1–222 (2008).
105. G. A. Lazkov, B. A. Sultanova, Checklist of vascular plants of Kyrgyzstan (Botanical Museum, Finnish Museum of Natural History, 2011).
106. Tropicos, Flora de Nicaragua.
107. Tropicos, Catálogo de las Plantas Vasculares de Bolivia.
108. Tropicos, Catalogue of the Vascular Plants of Ecuador.
109. Tropicos, Panama Checklist.
110. Chinese Virtual Herbarium, The Flora of China v. 5.0.
111. R. Bernal, S. R. Gradstein, M. Celis, Catálogo de plantas y líquenes de Colombia.
112. Jardim Botânico do Rio de Janeiro, Flora do Brasil 2020 em construção.
113. J. Danihelka, J. Chrtek, Z. Kaplan, Checklist of vascular plants of the Czech Republic. Preslia 84, 647–811 (2012).
114. T. Nikolić, Flora Croatica Database.
115. S. M. Kuzmenkova, Plants of Belarus (deposited 28 October 2015).
116. J. Thomas, Plant diversity of Saudi Arabia: Flora checklist.
117. A. W. A. Al Khulaidi, Flora of Yemen (2000).

118. Ministry of Environment, The national red list 2012 of Sri Lanka: Conservation status of the fauna and flora (2012).

119. T. Karlsson, M. Agestam, Checklist of Nordic vascular plants: Sweden Checklist.

120. P. Dimopoulos, T. Raus, E. Bergmeier, T. Constantinidis, G. Iatrou, S. Kokkini, A. Strid, D. Tzanoudakis, Vascular plants of Greece: An annotated checklist (Botanischer Garten und Botanisches Museum Berlin-Dahlem, Freie Universität Berlin; Hellenic Botanical Society, 2013).

121. Bundesamt für Naturschutz, Floraweb.

122. M. A. Hyde, B. T. Wursten, P. Ballings, M. Coates Palgrave, Flora of Botswana.

123. M. G. Bingham, A. Willemen, B. T. Wursten, P. Ballings, M. A. Hyde, Flora of Zambia.

124. M. A. Hyde, B. T. Wursten, P. Ballings, M. Coates Palgrave, Flora of Zimbabwe.

125. M. A. Hyde, B. T. Wursten, P. Ballings, M. Coates Palgrave, Flora of Mozambique.

126. M. A. Hyde, B. T. Wursten, P. Ballings, M. Coates Palgrave, Flora of Malawi.

127. M. A. Fischer, W. Adler, K. Oswald, Exkursionsflora für Österreich, Liechtenstein und Südtirol: Bestimmungsbuch für alle in der Republik Österreich, im Fürstentum Liechtenstein und in der Autonomen Provinz Bozen (Oberöstereichisches Landesmuseum, 2008).

128. C.-S. Chang, H. Kim, K. Chang, Provisional Checklist of the Vascular Plants for the Korea Peninsular Flora (KPF): Version 1.0.

129. VicFlora, Flora of Victoria.

130. G. Dauby, R. Zaiss, A. Blach-Overgaard, L. Catarino, T. Damen, V. Deblauwe, S. Dessein, J. Dransfield, V. Droissart, M. C. Duarte, H. Engledow, G. Fadeur, R. Figueira, R. E. Gereau, O. J. Hardy, D. J. Harris, J. de Heij, S. Janssens, Y. Klomberg, A. C. Ley, B. A. MacKinder, P. Meerts, J. L. van de Poel, B. Sonké, M. S. M. Sosef, T. Stévart, P. Stoffelen, J.-C. Svenning, P. Sepulchre, X. van der Burgt, J. J. Wieringa, T. L. P. Couvreur, RAINBIO: A mega-database of tropical African vascular plants distributions. PK 74, 1–18 (2016).

131. CONABIO, Sistema Nacional de Información sobre Biodiversidad.

132. INBIO, Lista de planta de Costa Rica: With updates by Eduardo Chacón.

133. Tropicos, Peru Checklist.

134. W.L.M. Tamis, R. van der Meijden, J. Runhaar, R. M. Bekker, W. A. Ozinga, B. Odé, I. Hoste, Standard List of the Flora of the Netherlands 2003. Gorteria 30, 101–195 (2004).

135. E. Buchwald, P. Wind, H. H. Bruun, P. F. Møller, R. Ejrnæs, H. E. Svart, Hvilke planter er hjemmehørende i Danmark? Jydsk Naturhistorisk Forening 118, 72–96 (2013).

136. I. M. Turner, A catalogue of the vascular plants of Malaya (Gardens' Bulletin, 1995).

137. Western Australian Herbarium, FloraBase - the Western Australian Flora.

138. Royal Botanic Gardens and Domain Trust, PlantNET - The NSW Plant Information Network System.

139. J. M. Lord, Patterns in floral traits and plant breeding systems on Southern Ocean Islands. AoB Plants 7, plv095 (2015).

140. S.-W. Breckle, I. C. Hedge, M. D. Rafiqpoor, Vascular plants of Afghanistan: An augmented checklist (Scientia Bonnensis, 2013).

141. G. Keppel, T. W. Gillespie, P. Ormerod, G. A. Fricker, Habitat diversity predicts orchid diversity in the tropical south-west Pacific. J Biogeogr 43, 2332–2342 (2016).

142. Landcare Research New Zealand, Ecological Traits of New Zealand Flora.

143. K. Yonekura, T. Kajita, BG Plants Japanese name and Scientific name Index (YList).

144. C. A. Backer, Bakhuizen van den Brink, RC, Flora of Java (Noordhoff, 1963).

145. Euro+Med, Euro+Med PlantBase - the information resource for Euro-Mediterranean plant diversity.

146. G. Zizka, Flowering plants of Easter Island. Palmarum hortus francofurtensis 3, 3–108 (1991).

147. M. D. Garroutte, Species Richness, Community Compositon and Species Distribution Patterns in Aleutian Plants: Thesis submitted in partial fulfillment of the requirements for the degree of Master of Science (2016).

148. C. Ulloa Ulloa, P. Acevedo-Rodríguez, S. Beck, M. J. Belgrano, R. Bernal, P. E. Berry, L. Brako, M. Celis, G. Davidse, R. C. Forzza, S. R. Gradstein, O. Hokche, B. León, S. León-Yánez, R. E. Magill, D. A. Neill, M. Nee, P. H. Raven, H. Stimmel, M. T. Strong, J. L. Villaseñor, J. L. Zarucchi, F. O. Zuloaga, P. M. Jørgensen, An integrated assessment of the vascular plant species of the Americas. Science (New York, N.Y.) 358, 1614–1617 (2017).

149. A. V. Yena, Prirodnaja flora krymskogo poluostrova: Spontaneous flora of the Crimean Peninsula (Orianda, 2012).

150. Tropicos, Catalogue of the Vascular Plants of Madagascar.

151. T. Muer, H. Sauerbier, F. Cabrera Calixto, Die Farn- und Blütenpflanzen der Kanarischen Inseln: Über 2.000 Pflanzenarten, mehr als 2.600 Fotos (Margraf Publishers, 2016).

152. M. Newman, A checklist of the vascular plants of Lao PDR (Royal Botanic Garden Edinburgh, 2007).

153. M. Jongbloed, G. R. Feulner, B. Böer, A. R. Western, The comprehensive guide to the wild flowers of the United Arab Emirates (Environmental Research and Wildlife Development Agency, 2003).

154. W. R. Sykes, Flora of the Cook Islands (National Tropical Botanical Garden, 2016).

155. R. Rodriguez, C. Marticorena, D. Alarcón, C. Baeza, L. Cavieres, V. L. Finot, N. Fuentes, A. Kiessling, M. Mihoc, A. Pauchard, E. Ruiz, P. Sanchez, A. Marticorena, Catálogo de las plantas vasculares de Chile. Gayana Bot. 75, 1–430 (2018).

156. P. L. Fall, T. D. Drezner, Vascular plant species of the Kingdom of Tonga by vegetation type, species origin, growth form, and dispersal mechanism. Ecology 101, e02902 (2020).

157. J. Hutchinson, J. M. Dalziel, R.W.J. Keay, N. Hepper, Flora of West Tropical Africa (Crown agents for overseas governments, 2014).

158. J. R. Press, M. J. Short, N. J. Turland, Flora of Madeira (Pelagic Publishing, 2016).

159. R. Cámara-Leret, D. G. Frodin, F. Adema, C. Anderson, M. S. Appelhans, G. Argent, S. Arias Guerrero, P. Ashton, W. J. Baker, A. S. Barfod, D. Barrington, R. Borosova, G. L. C. Bramley, M. Briggs, S. Buerki, D. Cahen, M. W. Callmander, M. Cheek, C.-W. Chen, B. J. Conn, M. J. E. Coode, I. Darbyshire, S. Dawson, J. Dransfield, C. Drinkell, B. Duyfjes, A. Ebihara, Z. Ezedin, L.-F. Fu, O. Gideon, D. Girmansyah, R. Govaerts, H. Fortune-Hopkins, G. Hassemer, A. Hay, C. D. Heatubun, D. J. N. Hind, P. Hoch, P. Homot, P. Hovenkamp, M. Hughes, M. Jebb, L. Jennings, T. Jimbo, M. Kessler, R. Kiew, S. Knapp, P. Lamei, M. Lehnert, G. P. Lewis, H. P. Linder, S. Lindsay, Y. W. Low, E. Lucas, J. P. Mancera, A. K. Monro, A. Moore, D. J. Middleton, H. Nagamasu, M. F. Newman, E. Nic Lughadha, P. H. A. Melo, D. J. Ohlsen, C. M. Pannell, B. Parris, L. Pearce, D. S. Penneys, L. R. Perrie, P. Petoe, A. D. Poulsen, G. T. Prance, J. P. Quakenbush, N. Raes, M. Rodda, Z. S. Rogers, A. Schuiteman, P. Schwartsburd, R. W. Scotland, M. P. Simmons, D. A. Simpson, P. Stevens, M. Sundue, W. Testo, A. Trias-Blasi, I. Turner, T. Utteridge, L. Walsingham, B. L. Webber, R. Wei, G. D. Weiblen, M. Weigend, P. Weston, W. de Wilde, P. Wilkie, C. M. Wilmot-Dear, H. P. Wilson, J. R. I. Wood, L.-B. Zhang, P. C. van Welzen, New Guinea has the world's richest island flora. Nature 584, 579–583 (2020).

160. Elizabeth Joyce, Kevin Thiele, Ferry Slik, Darren Crayn, E. Joyce, K. Thiele, F. Slik, D. Crayn, Checklist of the vascular flora of the Sunda-Sahul Convergence Zone. Pensoft Publishers 8, e51094 (2020).

161. BGCI, GlobalTreeSearch online database (version 1.4). Available at http://dx.doi.org/10.13140/RG.2.2.22578.84163.

162. I. G. Alsos, D. Ehrich, P. B. Eidesen, H. Solstad, K. B. Westergaard, P. Schönswetter, A. Tribsch, S. Birkeland, R. Elven, C. Brochmann, Long-distance plant dispersal to North Atlantic islands: colonization routes and founder effect. AoB Plants 7, plv036 (2015).

163. F. J. Burton, Red list assessment of Cayman Islands' native flora for legislation and conservation planning (2007).

164. S. J. Meades, S. G. Hay, L. Brouillet, Annotated checklist of the vascular plants of Newfoundland and Labrador (The Provincial Museum of Newfoundland and Labrador, St. John’s, NF, Canada., 2005).

165. V. Boullet, J. Hivert, L. D. B. Gigord, An Updated Account of the Vascular Flora of the Iles Eparses (Southwest Indian Ocean). Atoll Research Bulletin 614, 1–64 (2018).

166. T.-C. Huang, Flora of Taiwan, vol. I–VI (National Taiwan University, 1994).

167. R. R. Thaman, A. Tye, Flora of Kiritimati (Christmas) Atoll, Northern Line Islands, Republic of Kiribati. Atoll Research Bulletin 608, 1–73 (2015).

168. E. Freid, J. Francisco-Ortega, B. Jestrow, Endemic Seed Plants in the Bahamian Archipelago. Bot. Rev. 10.1007/s12229-014-9137-z, 1–27 (2014).

169. Virtual Biodiversity Museum of Cyprus, Flora of Cyprus - Endemic plants of Cyprus.

170. J. Francisco-Ortega, F.-G. Wang, Z.-S. Wang, F.-W. Xing, H. Liu, H. Xu, W.-X. Xu, Y.-B. Luo, X.-Q. Song, S. Gale, D. E. Boufford, M. Maunder, S.-Q. An, Endemic Seed Plant Species from Hainan Island: A Checklist. The Botanical Review 76, 295–345 (2010).

171. S. Mifsud, MaltaWildPlants.com - an online flora of Malta.

172. M. Rhind, Terrestrial Biozones: Endemic Floras.

173. C. M. S. Carrington, R. D. Edwards, G. A. Krupnick, Assessment of the Distribution of Seed Plants Endemic to the Lesser Antilles in Terms of Habitat, Elevation, and Conservation Status. Caribbean Naturalist Special Issue No. 2, 30–47 (2018).

174. R. Graveson, Plants of Saint Lucia: A Pictorial Flora of Wild and Cultivated Vascular Plants.

175. P. Jaramillo Díaz, A. Guézou, A. Mauchamp, A. Tye, CDF Checklist of Galapagos Flowering Plants: FCD Lista de especies de Plantas con flores Galápagos (Charles, 2018).

176. R. Govaerts, E. Nic Lughadha, N. Black, R. Turner, A. Paton, The World Checklist of Vascular Plants, a continuously updated resource for exploring global plant diversity. Scientific Data 8, 1 (2021).

177. D. Penneckamp Furniel, Flora vascular silvestre del Archipiélago Juan Fernández (Planeta de Papel Ediciones, 2018).

178. V. L. Komarov, B. K. Shishkin, E. G. Bobrov, Flora of the U.S.S.R (Israel Program for Scientific Translations, 1933-1964).

179. K. Stephens, Comparative floristic analysis of vegetation on the Dune Islands of South-East Queensland. The Proceedings of the Royal Society of Queensland 117, 141–180 (2011).

**SI References**


1. P. Weigelt, C. König, H. Kreft, GIFT – A global inventory of floras and traits for macroecology and biogeography. *J. Biogeogr.* **47**, 16–43 (2020).

2. R. Govaerts, E. Nic Lughadha, N. Black, R. Turner, A. Paton, The World Checklist of Vascular Plants, a continuously updated resource for exploring global plant diversity. *Sci. Data* **8**, 215 (2021).

3. G. Kier, *et al.*, A global assessment of endemism and species richness across island and mainland regions. *Proc. Natl. Acad. Sci.* **106**, 9322–9327 (2009).

4. B. Sandel, *et al.*, Current climate, isolation and history drive global patterns of tree phylogenetic endemism. *Glob. Ecol. Biogeogr.* **29**, 4–15 (2020).

5. I. R. McFadden, *et al.*, Temperature shapes opposing latitudinal gradients of plant taxonomic and phylogenetic β diversity. *Ecol. Lett.* **22**, 1126–1135 (2019).

6. A. Stein, K. Gerstner, H. Kreft, Environmental heterogeneity as a universal driver of species richness across taxa, biomes and spatial scales. *Ecol. Lett.* **17**, 866–880 (2014).

7. W. Jetz, C. Rahbek, R. K. Colwell, The coincidence of rarity and richness and the potential signature of history in centres of endemism. *Ecol. Lett.* **7**, 1180–1191 (2004).

8. G. G. Mittelbach, *et al.*, Evolution and the latitudinal diversity gradient: speciation, extinction and biogeography. *Ecol. Lett.* **10**, 315–331 (2007).

9. K. Rohde, Latitudinal gradients in species diversity: the search for the primary cause. *Oikos* **65**, 514–527 (1992).

10. G. C. Stevens, The latitudinal gradient in geographical range: how so many species coexist in the tropics. *Am. Nat.* **133**, 240–256 (1989).

11. B. J. Enquist, *et al.*, The commonness of rarity: Global and future distribution of rarity across land plants. *Sci. Adv.* **5**, eaaz0414 (2019).

12. R. Jansson, Global patterns in endemism explained by past climatic change. *Proc. R. Soc. Lond. B Biol. Sci.* **270**, 583–590 (2003).

13. B. Sandel, *et al.*, The influence of Late Quaternary climate-change velocity on species endemism. *Science* **334**, 660–664 (2011).

14. R. G. Gillespie, G. K. Roderick, Arthropods on islands: colonization, speciation, and conservation. *Annu. Rev. Entomol.* **47**, 595–632 (2002).

15. A. Antonelli, *et al.*, Geological and climatic influences on mountain biodiversity. *Nat. Geosci.* **11**, 718–725 (2018).

16. C. Rahbek, *et al.*, Building mountain biodiversity: Geological and evolutionary processes. *Science* **365**, 1114–1119 (2019).

17. K. D. Bennett, P. C. Tzedakis, K. J. Willis, Quaternary refugia of north European trees. *J. Biogeogr.* **18**, 103–115 (1991).

18. L. M. J. Dagallier, *et al.*, Cradles and museums of generic plant diversity across tropical Africa. *New Phytol.* **225**, 2196–2213 (2020).

19. A. S. Jump, C. Mátyás, J. Peñuelas, The altitude-for-latitude disparity in the range retractions of woody species. *Trends Ecol. Evol.* **24**, 694–701 (2009).
